# Genomic diversity of the African malaria vector *Anopheles funestus*

**DOI:** 10.1101/2024.12.14.628470

**Authors:** Marilou Boddé, Joachim Nwezeobi, Petra Korlević, Alex Makunin, Ousman Akone-Ella, Sonia Barasa, Mahamat Gadji, Lee Hart, Emmanuel W. Kaindoa, Katie Love, Eric R. Lucas, Ibra Lujumba, Mara Máquina, Sanjay Nagi, Joel O. Odero, Brian Polo, Claire Sangbakembi, Samuel Dadzie, Lizette L. Koekemoer, Dominic Kwiatkowski, Erica McAlister, Eric Ochomo, Fredros Okumu, Krijn Paaijmans, David P. Tchouassi, Charles S. Wondji, Diego Ayala, Richard Durbin, Alistair Miles, Mara K. N. Lawniczak

**Author notes:** These authors contributed equally (Marilou Boddé, Joachim Nwezeobi).

## Abstract

*Anopheles funestus* s.s. is a formidable human malaria vector across sub-Saharan Africa. To understand how the species is evolving, especially in response to malaria vector control, we sequenced 656 modern specimens (collected 2014-2018) and 45 historic specimens (collected 1927-1967) from 16 African countries. We find high levels of genetic variation with clear and stable continental patterns. Six segregating inversions might be involved in adaptation of local ecotypes. Strong recent signals of selection centred on canonical insecticide resistance genes are shared by multiple populations. A promising gene drive target in *An. gambiae* is highly conserved in *An. funestus*. This work represents a significant advance in our understanding of the genetic diversity and population structure of *An. funestus* and will enable smarter targeted malaria control.

## Main text

The mosquito species *Anopheles funestus* (Giles, 1900) has an extraordinary adaptive potential demonstrated by its vast geographic range that spans sub-Saharan Africa (*1*) (shaded region in Fig. 1a). Importantly, *An. funestus* is highly anthropophilic (*2*), has a significantly longer lifespan than other African malaria vectors (*3*), and in some areas is also reported to have extremely high *Plasmodium* infection rates (*4–6*). In much of eastern and southern Africa, the species is the major malaria vector (*7*). Like the vectors in the Gambiae Complex, the species is contributing to outdoor, early evening and late morning biting in response to the use of indoor-based interventions, including bed nets (*8–10*). Although it can be difficult to find as larvae, it may have an extended transmission season in some locations, probably due to a preference for larval habitats that persist in the dry months (*11*). *An. funestus* harbours several polymorphic chromosomal inversions that play an important role in adaptation (*12*). Similar to the three other major African human malaria vectors, which are all members of the Gambiae Complex, *An. funestus* exhibits insecticide resistance across its range (*13*) and in some locations, even higher tolerance to insecticides than *An. Gambiae* (*14*). Alleles that confer resistance can be shared across great distances or can differ locally, indicating a complicated mix of population connectivity and selective pressures (*15*). These characteristics, combined with its widespread distribution, make *An. funestus* a critical target for malaria control efforts. Substantial genomic data resources exist for the three major malaria vector species of the Gambiae Complex (*16*) and these have become foundational for the study and implementation of control efforts (*17*). Here, we establish a baseline understanding of genetic diversity, population structure, inversion frequencies, and historic and current selection in *An. funestus* across the African continent at a whole genome level.

**Fig. 1.**
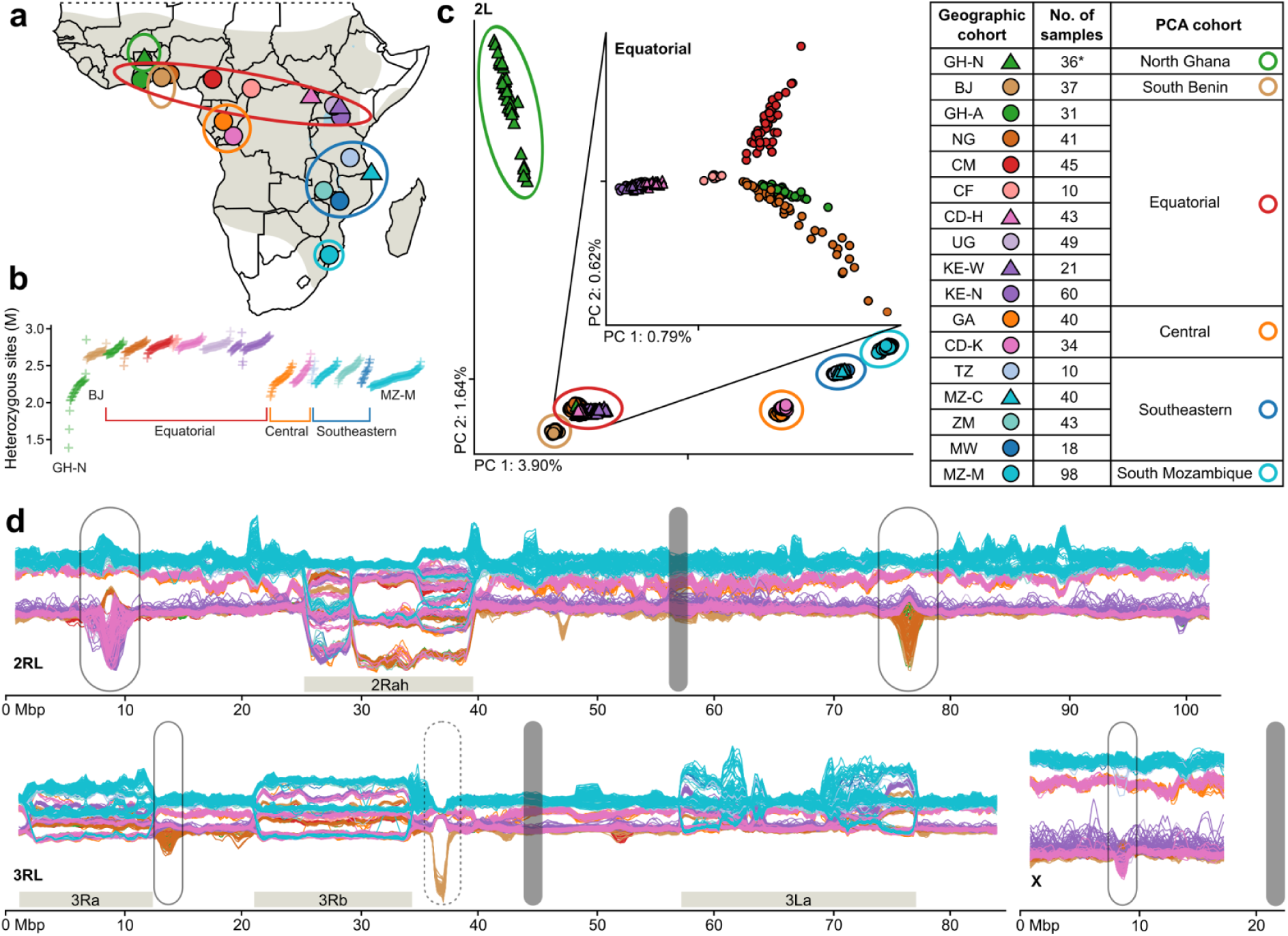
Population structure of 656 *Anopheles funestus* specimens collected across Africa. (**a**) Map showing sequenced individuals from sub-Saharan Africa grouped into 17 geographic cohorts based on their collection location (filled circular and triangular markers), and six PCA cohorts consisting of one or more geographic cohorts (open elliptical markers), with the grey shaded area showing the *An. funestus* range across the continent (adapted from Sinka et al. (*1*)). (*) One sample collected in GH-N clusters with Equatorial on the PCA and has therefore been removed in analyses using PCA-cohorts. (**b**) Number of heterozygous sites (in millions) per individual, with each cross representing one individual in subset_2 (females only, for subset_1 see fig. S2a) and ordered along the horizontal axis by geographic cohort (colour, same order as the legend) and number of heterozygous sites. (**c**) Projection along the first two principal components (PCs) computed on chromosome arm 2L, with the percentage of explained variance in the axis labels, and ticks denoting the 0 position along each axis. The PCA cohorts are indicated with open elliptical markers. The inset PC plot was computed only on samples from the Equatorial PCA cohort. (**d**) Sliding window PCA along the genome, each line corresponds to one individual, individuals from GH-N are excluded. The genome is split in overlapping windows of approximately 1 Mbp and 200 kbp steps, a PCA is performed for each window, PC1 values are plotted along the y-axis, and the window centres in genomic coordinates are plotted along the x-axis. (The corresponding plot displaying the PC2 values for the same windows is shown in fig. S5a). Inversion regions are indicated by horizontal grey bars, centromeres by vertical grey bars, lined ellipses indicate regions of divergence across several cohorts (Fig. 3, figs. S6-8), while the dotted ellipse showcases divergence specific to the South Benin PCA cohort (fig. S3c,d).

### Sample acquisition, sequencing, and genetic diversity

In 2017, we put out an open call to join the project through contributing wild *An. funestus* samples collected from 2014 onwards. We carried out short read sequencing at ∼35x coverage depth for over 800 wild caught specimens, of which 656 individuals from 13 African countries passed all quality controls (Fig. 1a, fig. S1a, and Methods). Sequencing reads were aligned to a 251 Mbp high quality chromosomal reference genome created from a wild caught individual from Gabon (*18*), and each successfully sequenced sample was assigned to a geographic cohort based on its original collection location (Fig. 1a, table S1). Using a static-cutoff (sc) site filter, 73 million (M) out of 162 M (45%) accessible sites are segregating among the sequenced samples (fig. S1d, Methods). Disregarding singletons, 49 M single nucleotide polymorphisms (SNPs) are present on two or more chromosomes, 17.3% of which have more than two alleles. For some analyses, we use a more stringent decision-tree (dt) site filter that results in 48 M out of 114 M (42%) segregating accessible sites, with 31 M present on two or more chromosomes, and 15.2% having more than two alleles. Genome-wide nucleotide diversity is between 1.4% and 1.7% in each geographic cohort (fig. S2c), with most individuals having between 2 and 3 million heterozygous sites, and higher diversity among equatorial individuals and cohorts (Fig. 1b and fig. S2a,c). Three mosquitoes from North Ghana have exceptionally low average heterozygosity (under 2 M variants) due to long runs of homozygosity (ROH, fig. S2b).

### Population structure

Principal Components Analysis (PCA) on variants from chromosome arm 2L (thought to have no common chromosomal inversions (*19*)), showed PC1 is correlated with latitude (Fig. 1c). Population structure is similar across other autosomal arms, but complicated by signals relating to segregating inversions (fig. S1e). We used the observed clusters in the PCA plot to define six PCA cohorts, five of which (North Ghana, South Benin, Central, Southeastern, and South Mozambique) contain relatively proximal geographic cohorts, whereas the sixth Equatorial cohort includes samples from seven countries spanning a 4,000 km range from Ghana to Kenya (Fig. 1a,c), suggesting a high degree of genetic connectivity across the equatorial region of Africa. Further PCA on the Equatorial cohort alone shows that the samples are clustered by geographic location (Fig. 1c, inset). Although the North Ghana and South Benin cohorts belong to *An. funestus* (fig. S3a) and are geographically close to the Equatorial cohort, they are genetically distinct (explored in more detail below). We also explored the mitochondrial genome for every individual, finding that the mitochondrial tree is largely consistent with the PCA cohorts (fig. S9e), although ten geographic cohorts have a small number of individuals falling in a separate mitochondrial lineage than the majority, and more than half of the South Mozambique samples are placed in ‘Lineage II’, which also contains other species from the Funestus Subgroup (*20*, *21*).

Samples from the Equatorial cohort display more negative Tajima’s D values than those from other PCA cohorts supporting an expanding population (fig. S2d). Pairwise fixation indices (F_ST_) and patterns of doubleton sharing support the PCA cohort groupings (fig. S2e,f). PCA was computed on common variants (minor allele frequency >0.01) that tend to be older, whilst doubleton sharing focuses on rare variants that tend to be more recent (*22*), suggesting that the observed structure does not differ considerably between historic and recent events. While further sampling across the continent is needed, these results indicate that there is one genetically connected population of *An. funestus* across the equator, while other populations both geographically proximal and distant to the Equatorial cohort are much more genetically isolated.

### Ecotypes and differentiated populations

Our understanding of *An. funestus* ecotypes has been enhanced through population genomic sequencing of Kiribina and Folonzo, which are considered chromosomal forms and were originally characterised by non-random association of inversion karyotypes (*23*, *24*). This recent work showed genome wide differentiation between these sympatric ecotypes within Burkina Faso. The continentally distributed Folonzo ecotype has varying inversion frequencies and a preference to breed in vegetative ditches and swamps, while the geographically restricted Kiribina ecotype has a nearly fixed standard karyotype and has adapted to breed in rice fields (*23*, *24*). Integrating these previously sequenced specimens into our dataset, we find that outside the inversion regions, Folonzo clusters with the Equatorial cohort and Kiribina is genetically distinct from Folonzo (fig. S3b), consistent with previous findings (*23*). However, two cohorts we investigate here that are geographically proximal to the equator – North Ghana and South Benin – are even more genetically distinct from Folonzo than Kiribina is.

The North Ghana cohort has reduced levels of diversity, an excess of ROHs, and high divergence from all other cohorts, including from GH-A, less than 400 km away, consistent with a recent population bottleneck (Fig. 1b, figs. S2, S3d). Comparing the two populations from Ghana reveals elevated F_ST_ in the genomic regions associated with the 2R and 3L inversions (fig. S3e) and indeed, North Ghana appears to be an outlier for its 2R karyotype in comparison to its nearest neighbours (fig. S4e). We were also unable to karyotype North Ghana for the 3La inversion, due to high divergence on the 3L arm particularly in the inversion region (fig. S3e,f). One individual from North Ghana actually clustered with the GH-A population. Together, these karyotypic anomalies alongside a potentially sympatric population that looks to be Folonzo hint that the North Ghana cohort may be another ecotype, but further sampling will be needed to confirm this hypothesis.

The South Benin (BJ) cohort has similar levels of variation as the Equatorial cohort but is clearly differentiated in the PCA (Fig. 1c). To further explore this differentiation, we ran sliding window PCAs that result in a view of population structure along the genome (Fig. 1d, doi.org/10.5281/zenodo.13993020 for interactive plots; North Ghana was excluded from these analyses due to excessive ROH). While BJ follows the Equatorial cohort for most of the genome, it is highly differentiated from all other geographic cohorts in one genomic region (3R: 37Mb). F_ST_ between BJ and the neighbouring GH-A shows the strongest differentiation between the two populations falls between two genes characterised as ‘semaphorin-2A-like’ (AFUN2_001563, AFUN2_006509) (fig. S3c). We do not find any clear nonsynonymous differentiation between these populations and it may be that the divergence is regulatory in nature or that the reference genome is inadequately representing the sequences of these individuals in this genomic region (table S3). Populations from Benin are exceptionally resistant to DDT (0% mortality after one hour of exposure) (*25*), but there is currently no evidence in the literature to support a role in DDT resistance for semaphorin-2As, which are secreted or transmembrane signalling proteins; instead they have been implicated in nerve development, limb development, and olfaction (*26*, *27*). Strong divergence could be caused by differential regulation of genes involved in DDT resistance, assortative mating, or habitat adaptation, but genome accessibility drops off as F_ST_ increases and the cohort is too diverged from the reference genome in this genomic region to speculate further.

BJ has inversion frequencies that are also strongly contrasting with its close geographic neighbours: it is fixed standard for all inversions except 3La (Fig. 2, fig. S4e). This is reminiscent of the Kiribina ecotype, which is (nearly) fixed standard for 2Ra, 3Ra and 3Rb (*23*, *24*), however, BJ is clearly differentiated from Kiribina (fig. S3b). Furthermore, in this sample set, we find no evidence of a Folonzo-like population in South Benin, so it is unclear whether BJ is sympatric with Folonzo or is the only form present in the region.

**Fig. 2.**
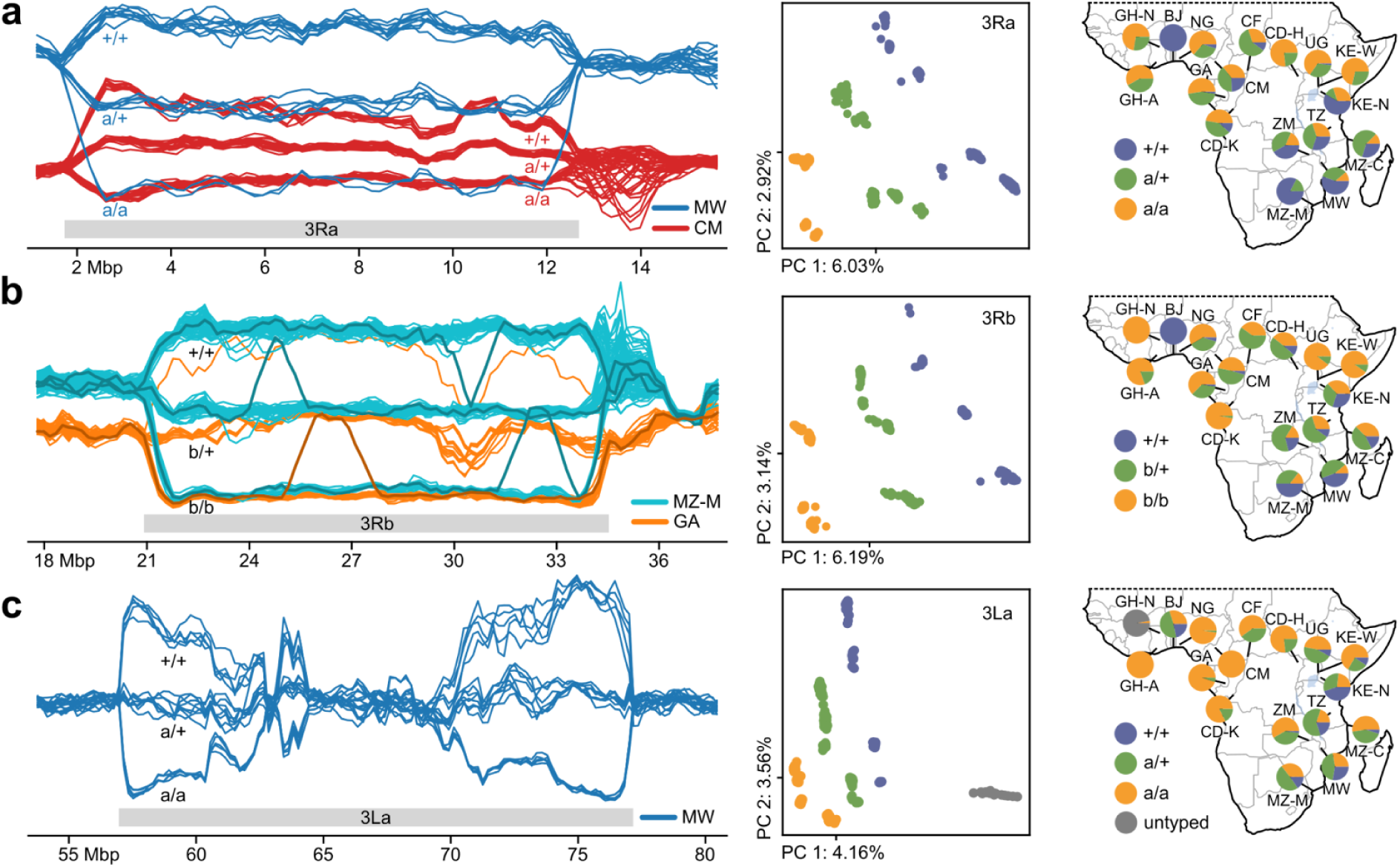
Segregating inversions on chromosome 3. Sliding window PCAs (left panels) were computed on all samples except North Ghana. For visualisation purposes, only a subset of samples is shown in the sliding window PCAs. (**a**) (left) Sliding window PCA showing CM (red) and MW (blue) samples on the 3Ra inversion region; window centres in Mbp along the x-axis, PC1 values for each individual along the y-axis. The different karyotypes are visible as three different horizontal trajectories, from top to bottom 3R+/+, 3Ra/+, and 3Ra/a. All three karyotypes are present in both cohorts, and because PC1 captures a signal that is a combination of the inversion and geographic population structure, the upper and middle trajectories (3R+/+ and 3Ra/+, respectively) do not overlap for the different cohorts, see fig. S5d. (middle) Projection along the first two principal components computed on the entire inversion region using all samples; samples are coloured by inversion karyotype. (right) Map of karyotype frequencies per geographic cohort. (**b**) (left) Sliding window PCA on the 3Rb inversion region displaying GA (orange) and MZ-M (cyan). Several samples change trajectory for part of the inversion region (indicated by thicker, darker lines); these samples appear to be double recombinants that locally exhibit a different karyotype than they have for the rest of the inversion (see fig. S5e, Supplementary Information). (middle and right) as in (a). (**c**) Sliding window PCA displaying MW on the 3La inversion region shows a strong separation of karyotypes near the breakpoints, but the signal decays towards the inversion centre (see fig. S5b). (middle and right) as in (a).

### Inversions

Chromosomal inversions can capture coadapted alleles and protect them from recombination (*28*). In the sliding window PCA, we find five large segregating inversions that drive population structure in their respective genomic regions, and after determining inversion breakpoints we identified these inversions as 2Ra, 2Rh, 3Ra, 3Rb, and 3La (Fig. 1d, fig. S1b, Supplementary text, table S2). We performed *in silico* karyotyping for all samples and all inversions using sliding window and aggregated PCA on the inversion region (Fig. 2, fig. S4, table S1). Inversion frequencies vary greatly across the continent and previous studies have linked inversions to behaviour and adaptation (*12*, *29*, *30*). Although Hardy-Weinberg equilibrium (HWE) of inversion karyotypes is typically satisfied within PCA cohorts, there are a few exceptions where samples collected in different seasons do not satisfy HWE when grouped together (Supplementary text). Changes in inversion frequencies during the wet and dry seasons have previously been reported for *An. funestus* (*31*) as well as the Gambiae Complex (*32*). The inversions also differ in heterozygosity of each homokaryotype: 2Rh/h has lower heterozygosity than 2R+^h^/+^h^, hinting that the standard orientation is likely ancestral (fig. S5d).

The sliding window PCA enables the investigation of local structure within the inversions. In the genomic regions where large inversions are segregating, PC1 captures a compound signal of inversion karyotype and population structure (Figs. 1d, 2, fig. S4-5). Size, age, recombination rates, and demographic history of the inversion can all affect the relative strength of the inversion and population structure signals, and the observed amount of genetic variation in the different inversion orientations (*33*). While for most inversions the relative strength of these two signals is constant for the entire inversion (e.g. Fig. 2a), for 3La the inversion signal decays in the middle (Fig. 2c and fig. S5b). This distinct pattern of 3La in comparison to the other inversions may have been driven by more frequent double recombination due to its large size or due to its age. Sliding window PCA reveals individuals with inversion karyotypes that are the product of double recombination in 3Rb and 3La (Fig. 2b, fig. S5e,f). Remarkably, these double recombinants can be seen in the sliding window plots where individuals change karyotype trajectories for a part of the inversion region (Fig. 2b). Events like these move alleles from one karyotype into the other, and are important for the spread of variants (*34*).

Consistent with Sharakhov *et al.* (*19*), we observe several overlapping inversions on chromosome arm 2R: 2Ra and 2Rh, which occur in many cohorts, and 2Rt, which occurs only in the CM and CD-H cohorts (Fig. 1d, fig. S4). There are six possible combined karyotypes for 2Rah, all resulting in unique trajectories in the sliding window PCA (fig. S4b,d). The relatively rare 2Rt inversion results in two additional states in the region where it overlaps 2Ra (fig. S4c,d). We believe that the 2Rt inversion only occurs in the heterozygous state in this dataset, firstly because that results in a consistent interpretation of the sliding window PCA trajectories of the combined 2Raht karyotypes, and secondly because the samples carrying 2Rt never interpolate between two groups of homozygotes, but drive a PC by themselves. Assuming that this dataset does not contain individuals homozygous for the 2Rt inversion, both CM and CD-H are in HWE.

### Signals of selection

To explore signals of selection, we computed H12 (*35*), a statistic measuring haplotype homozygosity along the genome (Fig. 3a, Methods). H12 quantifies excess homozygosity indicative of hard or soft selective sweeps. We identified four regions that are putatively under very strong selection (H12 > 0.4) in at least two geographic cohorts (Fig. 3a, grey boxes); notably, these regions are also visible as peaks in the sliding window PCA (Fig. 1d, circled regions). All four H12 peaks are centred on known insecticide resistance genes that are important in many insect species (*Gste2, Gaba, Cyp6p, Cyp9k1*) (*36*).

**Fig. 3.**
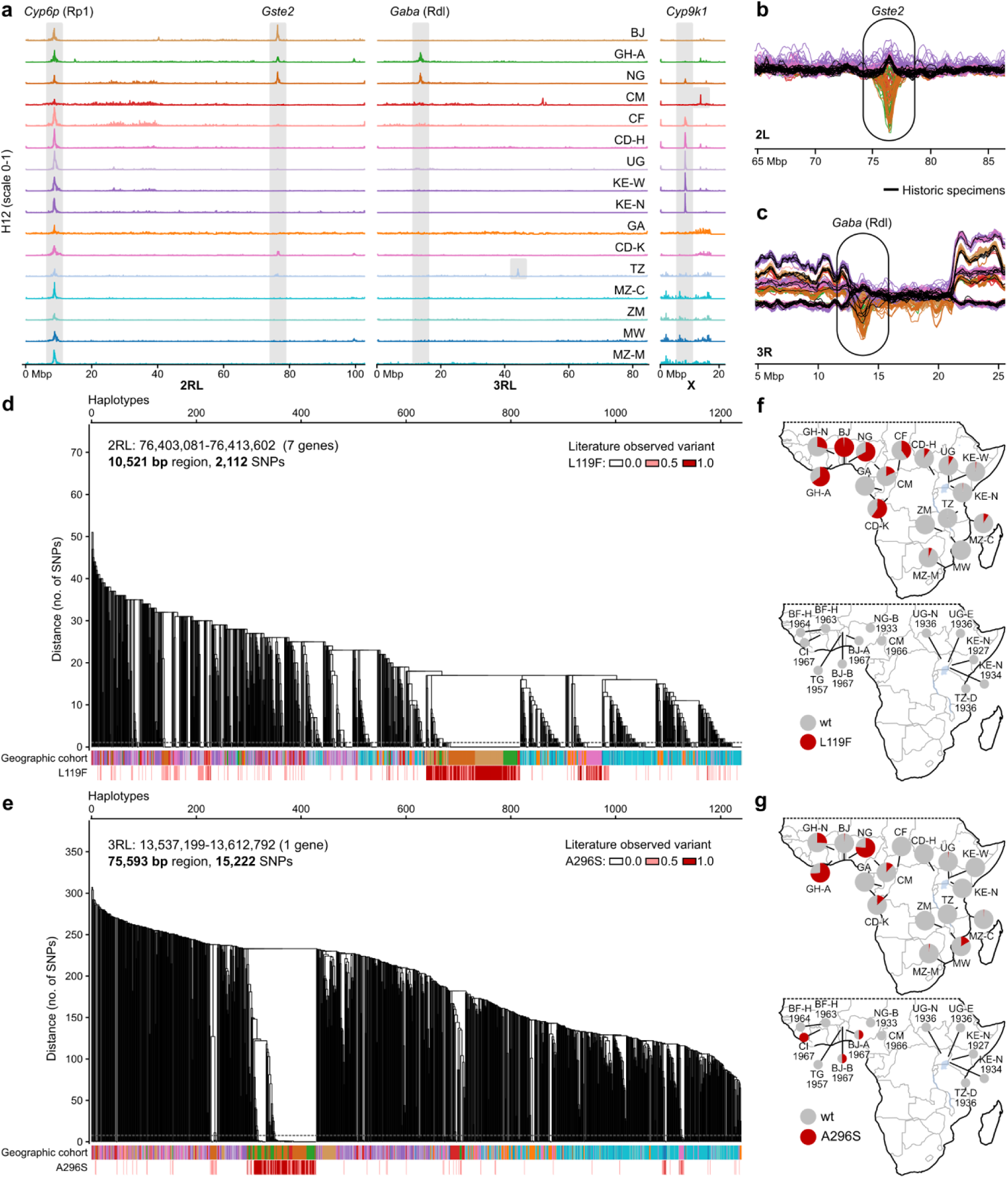
Selection sweeps and known insecticide resistance variants in *Gste2* and *Gaba*. (**a**) Genome-wide H12 scans for signals of recent selection. H12 values range from 0 to 1, with higher values indicating excessive haplotype sharing, which is a signature of recent selection. North Ghana was excluded due to elevated runs of homozygosity, which adds noise and makes H12 values unreliable for detecting genuine selection signals. The y-axis runs from 0 to 1 for each cohort, the x-axis shows positions along the genome. Peaks of H12 values ≥0.4 are highlighted with a grey vertical bar; for peaks where only one cohort reaches this threshold, the bar is restricted to this cohort. (**b**) Sliding window PCA computed on modern samples and historic samples combined, excluding North Ghana (fig. S9d). For visualisation purposes, only present-day Equatorial and South Benin cohorts and the historic Equatorial cohort are shown. Here displaying a 20 Mbp region around the *Gste2* gene. Present-day samples show a peak in the *Gste2* region, which is likely a signal of selection, while historic samples (shown in black) do not exhibit this signal. (**c**) Sliding window PCA computed and samples subsetted as in (b), here showing a 20 Mbp region around the *Gaba* gene. Several historic samples follow the peak seen in present-day samples at the *Gaba* locus. (**d**) Haplotype clustering within a region containing seven *Gste* genes. The dendrogram is obtained by hierarchical clustering of phased haplotypes, and used to define haplotype clusters as groups of haplotypes with SNP divergence below 0.0005 (cutoff indicated as dashed horizontal line on the dendrogram). The first bar below the dendrogram shows the population of origin for each haplotype, the second bar shows the genotype for the known *Gste2* L119F mutation (note that this mutation was filtered out before haplotype phasing, so each haplotype is coloured by the genotype of the individual it belongs to; fig. S6a, table S3). (**e**) Haplotype clustering within the *Gaba* gene. Same structure as in panel (d) with the red bar showing the genotype for the rdl A296S variant (fig. S6b, table S3). (**f**) Maps showcasing the frequency of the L119F mutation in present-day (above) and historic (below) sample sets. (**g**) Maps showcasing the frequency of the rdl A296S mutation in present-day (above) and historic (below) sample sets.

The H12 peak on chromosome arm 2L, observed in BJ, GH-A, NG, and CD-K, is centred on a cluster of seven glutathione S-transferase (*Gste*) genes. The non-synonymous L119F mutation in *Gste2* is important in both DDT and pyrethroid resistance in *An. funestus* (*25*). We observe this mutation across the continent, at low frequencies in the east, and at particularly high frequencies in the aforementioned cohorts, notably nearly fixed in BJ (Fig. 3a,d,f, table S3). Most individuals from BJ, GH-A, and NG with the L119F mutation share the same haplotype in the *Gste* region, while in CD-K the L119F mutation is found on a different haplotypic background (Fig. 3d, fig. S6a), suggesting two distinct selective sweeps, rather than spread of a single resistance mutation.

The H12 peak on chromosome arm 3R, in GH-A and NG, is centred on the gamma-aminobutyric acid receptor subunit beta (*Gaba*) gene (Fig. 3a, table S3). In many species, a mutation at one amino acid residue in *Gaba* confers dieldrin resistance, hence being named the resistance to dieldrin (rdl) mutation (*37*). The A296S rdl mutation in *An. funestus* has previously been detected at high frequency in Central and Western Africa (*38*). The same A296S mutation is found here in ten geographic cohorts, predominantly on two distinct swept haplotypic backgrounds that occur sympatrically in GH-A and NG (Fig. 3e,g), but also spread throughout the continent at low frequency, discussed further below.

The other strong H12 peaks occurring in multiple cohorts include two other well studied insecticide resistance loci, *rp1* and *Cyp9k1*. There are also strong H12 peaks that are only found in a single cohort (TZ, CM) including a clear sweep on the well studied knock down resistance (kdr) mutation at the voltage gated sodium channel (*vgsc*) locus only in the Tanzanian samples – these samples were collected from a region that has a DDT stockpile that was leaking into the environment for decades (*39*). These loci are all discussed in the Supplementary text and figs. S7-8.

It is clear that some of the strongest selective pressures observed in this species are driven by insecticides. Widespread use of synthetic insecticides began in Africa in the late 1940s with the advent of DDT (*40*, *41*). To begin to explore both when and where the mutations we observe above arose, we sequenced 75 historic mosquitoes labelled as *An. funestus* from museum collections spanning 1927-1973 (fig. S9, table S4, Methods) (*42*). Based on mitochondrial and nuclear sequencing data, only 45 of these (spanning 1927-1967) were actually *An. funestus* s.s. (table S4). These 45 samples originated from nine countries and clustered together with the Equatorial and Southeastern PCA cohorts, which is expected based on their origins (fig. S9c). We integrated the historic data into the sliding window PCA, which showed that most of the genome in historic samples followed their present-day cohorts well, suggesting that population structure has been relatively stable over the past century (fig. S9d, see doi.org/10.5281/zenodo.13993020 for interactive plots). Intriguingly, nine historic samples follow the trajectory of South Benin in the divergence peak in chromosome arm 3R, and the alleles observed in the historic samples are highly similar to those found in modern day South Benin (fig. S10). In South Benin, there is exceptionally strong DDT resistance (*43*) but it remains to be determined what this allele, which is now found only in South Benin, was associated with in these historic populations.

Emerging DDT resistance was phenotypically detected in *An. funestus* in the late 1950s (*44*), however, we do not find any evidence of either L119F in *Gste* or kdr in *vgsc* that are associated with DDT resistance today (table S4). We suspect that if the kdr mutation conferred resistance to DDT 60 years ago, it would have been rapidly reselected when pyrethroid treated bed nets were distributed at scale. Historic genomes also do not have evidence of insecticide resistance mutations at *rp1* or *Cyp9k1* (table S4), suggesting that selection pressures on these genes are more recent. However, we find that six individuals from the 1960s, the time when dieldrin resistance in *An. funestus* was first reported (*44*, *45*), follow the PC1 trajectory of present-day resistant populations that carry the rdl A296S mutation (Fig. 3b, fig. S9d) and indeed each of those individuals carry the mutation in a heterozygous or homozygous state. *Gaba* might act as a secondary target for pyrethroids (*46*), which could explain the persistence of rdl in modern populations. Alternatively, or additionally, dieldrin, like DDT, is a persistent organic pollutant and also has stockpiles across Africa. Perhaps leaking dieldrin stockpiles continue to exert selective pressure on modern populations or perhaps unregulated usage continues in some areas. Undoubtedly, these long lasting insecticides that are stored in poor conditions in many locations across Africa are complicating resistance management and vector control (*47*).

Today, while we observe that both the L119F mutation in *Gste2* and the rdl mutation in *Gaba* are on haplotypes undergoing selective sweeps in west and central Africa, they are also found scattered across the continent at lower frequencies (Fig. 3d-g). This may hint that historic selection for these mutations was relaxed during the period of the 1970s-1990s when DDT and dieldrin usage declined, but these mutations may have remained at low frequencies, preadapting the species to rapidly evolve resistance to pyrethroids when those came into common usage in the 2000s.

### Gene drive

The use of gene drive to eliminate or modify malaria vector species holds great promise as a targeted malaria vector control strategy (*48*). Knowledge of genetic variation and structure throughout the species range is important to both assess the likelihood of geographical spread and to identify candidate targets for CRISPR-Cas9 gene drive, given that polymorphism within target sites affects gene drive efficacy (*16*). Suppression based gene drive targeting the doublesex (*dsx*) gene has already been developed and successfully tested in the laboratory and large cage setting for *An. gambiae* (*49, 50*). Approaches developed for *An. gambiae* should be technically sound for *An. funestus*, but population structure and levels of genetic diversity will differ between these species.

Here we evaluated potential target sites within coding sequence for *An. funestus*. Gene drive targets are identified as 20 bp sequences located entirely within coding sequence, containing the protospacer adjacent motif (PAM). In total, we identified 2,349,313 target sites in 11,086 genes based on the reference genome. However, only 30,459 sites in 3,927 genes remained after excluding targets with variation in any subset_2 individual. In comparing target site availability in *An. funestus* to *An. gambiae,* we find that *An. gambiae* has more available targets based on the reference genome alone (2,718,188), but the decay in number of targets as we consider variation in cumulatively more individuals is similar to *An. funestus* (Fig. 4a, Supplementary text). We also specifically explored the *dsx* gene drive target region in *An. funestus*, finding that the reference genomes (AgamP4 and AfunGA1) only differ by one base pair (Fig. 4b), and only a single SNP is found segregating in two individuals among all the *An. funestus* we sequence here and this SNP is the *An. gambiae* reference allele. Altogether, this bodes well for using the same population suppression gene drive approach in *An. funestus* as is underway for *An. gambiae*.

**Fig. 4.**
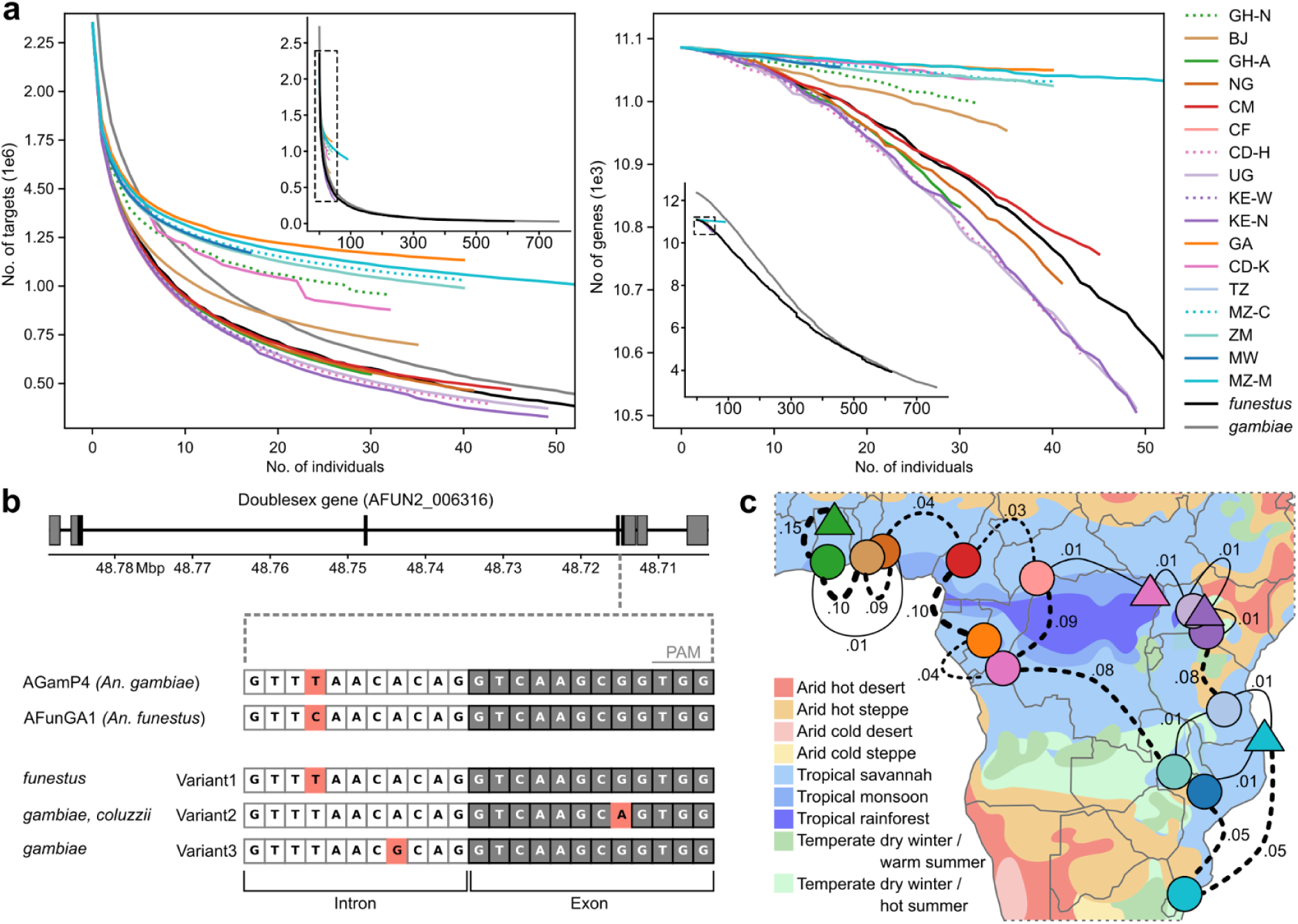
Challenges for vector control. (**a**) Cumulative number of gene drive targets (left) and number of genes containing gene drive targets (right) per geographic cohort as well as all subset_2 individuals (*funestus*) and *An. gambiae* and *An. coluzzii* individuals from Ag1000G phase 1 (*16*) (*gambiae*). The insets are zoomed out versions of the main plot, showing the results for the fully explored datasets; the areas of the main plots are indicated as dashed boxes in the insets. (**b**) Sequence variation in the *dsx* gene drive target on chromosome arm 2R. The *dsx* gene drive target is located at the boundary of intron 4 and exon 5 of the female-specific isoform. At the top, the seven exons of the female-specific isoform are depicted as boxes on the AGamP4 reference genome; coding sequence is shown in black, untranslated exonic regions (UTR) in grey. The target sequence is shown below, nucleotides in white boxes are located in intron 4, nucleotides in grey boxes are located in exon 5; the last three nucleotides constitute the PAM. The AGamP4 (*An. gambiae*) and AFunGA1 (*An. funestus*) reference genomes differ at the fourth base of the target sequence (highlighted in pink) in the intronic region. Variant 1 is found at very low frequencies in two cohorts from this study, Variant 2 is found at low to intermediate frequencies in four cohorts of *An. gambiae* and *An. coluzzii* and Variant 3 is found at very low frequency in one cohort of *An. gambiae* (see Supplementary Information). (d) Map showing fixation indices (F_ST_) for neighbouring populations. F_ST_ is calculated on all accessible sites on the 2L chromosome arm. F_ST_ > 0.01 lines are dotted and become thicker the higher the F_ST_. The background of the map is coloured by the Köppen-Geiger climate classification, adapted from Beck *et al.* (*51*).

## Discussion

Even if the Gambiae Complex disappeared today, malaria will still rage through Africa until *An. funestus* is also effectively targeted. The greater understanding of the high levels of genetic diversity and the complex population structure of *An. funestus* presented here will underpin smarter surveillance and targeted vector control (*3*, *52*, *53*). More than 4,000 kilometres separates the sampling sites of the Equatorial cohort (from Ghana Ashanti to Western Kenya), yet populations from across that range are genetically connected, while much geographically closer populations like South Benin and North Ghana are genetically distinct (Fig. 4c). Some of this structure may originate from geographic discontinuities like the Congo basin rainforest and the Rift valley, and some may originate from differences in climate and rainfall (*51*) (Fig. 4c). Small *et al.* (*23*) predicted that other ecotypes are likely to be found as genomic investigations into the species proceed (*54*, *55*), and we argue that both South Benin and North Ghana may provide two more examples. The identification of these ecotypes highlights the complex population structure of *An. funestus* and the need for locally tailored control strategies.

A comprehensive understanding of gene flow within and between each of the major malaria vector species is critical for implementing effective malaria control, whether that be through gene drive release or successful insecticide use and resistance management. The high genetic diversity and complex population structure of *An. funestus* suggests that a one-size-fits-all approach to vector control may be ineffective. The identification of strong selective sweeps centred on known insecticide resistance genes across multiple populations highlights the ongoing challenge of insecticide resistance. Notably, the same mutations tend to confer resistance and be under selection in a wide range of insects (*36*), and in both *An. funestus* and *An. gambiae* (*56*) these mutations sometimes occur on multiple distinct haplotypic backgrounds suggesting populations may not be mutation limited. This convergent evolution of resistance mechanisms highlights the adaptability of these vector populations and underscores the necessity of tailored interventions that consider local genetic backgrounds and resistance profiles (*57*). Insecticide resistance in *An. funestus* results from a complex interplay between convergent evolution, sharing of resistance alleles between populations and changing selective pressures in space and time, and calls for the continued monitoring of resistance alleles alongside the development and deployment of novel insecticides or alternative control strategies (*58*). Population suppression or modification gene drives are a promising alternative to insecticide-based control and we identified a comparable number of candidate gene drive targets in *An. funestus* as in *An. gambiae* and *An. coluzzii*.

This work represents the first step in generating a foundational genomic understanding of *An. funestus* akin to what we have for the other major malaria vector species in Africa (*16*, *59*). Population genomic data resources are increasingly critical for informing vector control and are needed to underpin both technical strategies and global health policies. Further studies will need to focus on exploring the full range of the species given the complex pattern of population connectivity, and should integrate phenotypic data to understand the functional significance of observed genotypes. Temporal sampling is a key component of vector surveillance in order to comprehend the dynamics of genetic changes, e.g. frequency changes of resistance alleles, seasonal changes in inversion frequencies, and the range and persistence of ecotypes. Beyond increased spatiotemporal sampling, long read sequencing of ecotypes and resistance alleles will enhance our understanding of gene flow in this important malaria vector.

## Acknowledgments

We thank Katharina von Wyschetzki and Sonia Goncalves for assistance in sequencing submissions and Sanger’s Scientific Operation Teams for carrying out all library generation and sequencing on data presented here. We also express our gratitude to the entomologists who decades ago collected and identified the *Anopheles* specimens used in this study: Adam, Brunhes, Buxton, Garnham, Hamon, Leeson, Lewis, Pichon, Rickenbach, Stymes, IRD collections, and all supporting fieldwork and collection assistants whose names did not fit on the specimen labels.

## Funding

Wellcome Trust Quinquennial awards 206194 and 220540/Z/20/A(MKNL, JN, AMa, PK, and sequencing and also supported DK, who died in 2023)

Wellcome 4 year PhD studentship RG92770 (MB)

Bill & Melinda Gates Foundation INV-001927 (AMi, LH) Wellcome award WT207492 (RD)

National Research Foundation of South Africa SRUG2203311457 (LK)

Gates Foundation Grand Challenges Grant No. INV024969 and Investment Grant OPP1210769 (BP, EO)

Wellcome Trust International Intermediate fellowship 222005/Z/20/Z (DPT) Wellcome Trust Senior Research fellowship 217188/Z/19/Z (CSW)

ANR, France ANR-18-CE35-0002-01–WILDING (DA)

Bill and Melinda Gates Foundation and Obra Social “la Caixa” Partnership for the Elimination of Malaria in Southern Mozambique INV-008483 (KP, MM)UK Medical Research Council MR/T001070/1 (ERL, SJN, awarded to Martin Donnelly)

US National Institute of Health R01AI116811 (ERL, SJN, awarded to Martin Donnelly)

## Author contributions

MKNL conceived of and coordinated the study. OA, MG, EK, MM, CS, SD, LLK, EM, EO, FO, KP, DPT, CSW, and DA provided samples and secured funding to do so. PK and KL carried out DNA extractions. DK and MKNL secured the funding to support the sequencing and analysis. JN led the data curation with contributions from LH and AMi. MB led the data analysis with major contributions from JN and PK and minor contributions from AMa, SB, EL, IL, SN, JOO, and RD. MB and MKNL wrote the manuscript with major contributions from PK and JN and minor contributions from all co-authors. DA, RD, AMi, and MKNL supervised the lead authors.

## Data and materials availability

ENA accession IDs for each sample can be found in table S1 (modern) and table S4 (historic). All metadata associated with each sample is in the same tables. These data are released as part of the ENA BioProject PRJEB2141, the Vector Observatory Project. Processed data can be accessed through the malariagen_data python package at https://malariagen.github.io/vector-data/af1/af1.0.html. Analysis code will be available at <u>github.com/malariagen/funestus_popgen_ms</u> prior to publication.

## Supplementary Materials

### Materials and Methods

#### Population sampling

In 2017, we circulated an open call to vector biologists working in Africa to establish a baseline understanding of genomic diversity and population structure in *Anopheles funestus.* We received mosquito carcases or DNA collected from 13 countries between 2014 and 2018 (table S1, Fig. 1a). Specimens were collected indoors and outdoors using a variety of methods (human landing catch, pyrethroid spraying, manual aspiration, CDC light traps, and larval dipping followed by rearing to adulthood) and most were stored on silica gel after collection. Specimens were morphologically identified by collectors as *An. funestus*. The majority of specimens are females and comprise whole mosquitoes, but the dataset includes a small number of males and partial specimens (e.g. head/thorax or abdomen only). No phenotypic characterisation (e.g. bioassay outcomes, inversion karyotypes) was carried out on the specimens.

Additionally, a total of 75 dry pinned or ethanol stored historic specimens were selected from the anopheline collections at the London Natural History Museum (NHMUK) and Institut de Recherche pour le Développement in Montpellier (IRDFR) (table S4). These samples were collected from 10 countries between 1927 and 1973 (fig. S9c). Most samples were labelled as *An. funestus*, however 30 turned out to be *An. leesoni*, *An. rivulorum,* and another unknown Rivulorum Subgroup species, as observed from mitochondrial DNA ML tree clustering with previously published full *Anopheles* mitogenomes (fig. S9e).

#### Whole genome sequencing

Samples that arrived at the Wellcome Sanger Institute as mosquitoes were typically non-destructively extracted using Buffer C (*42*) and the lysates were purified using Qiagen MinElute kits (table S1). A small number of samples were extracted with Buffers A or G (*42*). Samples provided by contributors as DNA had been previously extracted in a variety of ways including CTAB and Qiagen DNeasy kits. All DNAs were quantified using Quant-iT picogreen dsDNA assays (ThermoFisher Scientific) following manufacturer protocols. Every DNA extract was subjected to a species-diagnostic PCR (*60*) and the majority showed the expected band size for *An. funestus.* A small number of samples were sequenced despite unclear results for the band-size based assay, but typically did not pass quality control (QC) as they turned out to be different species based on mitochondrial genome analysis (figs. S1a, S9e). Samples that had at least 70 ng of DNA were submitted for standard library preparation, which included shearing to 450 bp using a Covaris LE220 instrument, purification by SPRIselect Beads on an Agilent Bravo WorkStation, library construction using a custom protocol for the NEBNext Ultra II DNA Library Prep Kit for Illumina on an Agilent Bravo Workstation, tagging with KAPA HiFi HotStart ReadyMix and custom Integrated DNA Technologies (IDT) primers with Illumina UDI 1-96 barcodes, library quantification by qPCR, and finally pooling libraries in equimolar amounts before sequencing (*61*). For more than 200 samples with < 30 ng of DNA, libraries were prepared using an established low input method designed for laser capture microdissection that uses the NEBNextUltra II Fragmentase System (*62*). If samples had between 30 and 70 ng of DNA, they went for one of these two library preparation methods (table S1). Each pool, containing on average 30 individual samples, was sequenced on three lanes of an Illumina HiSeq X10 platform using PE150 kits, aiming for 30x coverage per individual.

Historic samples were extracted and sequenced following a previously described method (*42*). Briefly, samples were extracted with the same type of Buffer (mostly C, some A and G) but with shorter incubation times, and double-stranded library preparation was performed using the same NEBNext Ultra II DNA Library Prep Kit for Illumina with modifications to retrieve short and damaged DNA fragments. Libraries were then sequenced on an Illumina HiSeq or NovaSeq instrument using PE75 and PE150 kits.

#### Sequence data processing and variant calling

Alignment and genotyping was performed with the MalariaGEN alignment and genotyping pipelines with their default parameters (*63*, *64*). For each sample, sequencing reads were aligned to the AfunGA1 reference genome (*18*) using bwa mem v0.7.15 (*65*) and all alignments were post-processed and combined across lanes with samtools v1.4.1 (*66*) and Picard v2.9.2 (*67*). PCR duplicates were marked with the biobambam v2.0.73 (*68*) `bammarkduplicates` command. Reads were realigned around indels with GATK v3.7.0 (*69*) IndelRealigner, with target intervals generated for each sample separately. Variant calling was performed using GATK v3.7-0 (*69*) UnifiedGenotyper with all possible substitutions across all non-N sites of the reference genome marked as `--alleles`. Resulting VCF files were converted to a zarr format using the scikit-allel v1.2.1 (*70*) `vcf_to_zarr` function. Links to raw sequenced library fasta files, mapped BAM files, and both variant call files (VCF, zarr) per sample are listed in table S1.

Historic samples were preprocessed with the ancient DNA pipeline EAGER v2.3.5 (*71*), which performs adapter trimming, merging of paired-end reads if there is a ≥11 bp overlap, alignment with bwa mem v0.7.17 (*65*), removal of PCR duplicates and unmapped reads. Variant calls are generated as above. All libraries show characteristics of ancient DNA, such as C>T 5’ and G>A 3’ substitutions and short reads without prior shearing, highlighting we are indeed generating data from old mosquito DNA (table S4).

##### Sample and population quality control

All 838 samples underwent quality control (QC), which assessed median coverage, fraction of genome covered, divergence from the reference genome, estimated contamination percentage, sex determination, and replication likelihood. Any sample that failed to meet our specified parameter thresholds was excluded from further analysis. Sample failures appeared to be random, irrespective of DNA extraction type or library preparation method (table S1). The thresholds we used are: a minimum of 10x median coverage, at least 85% of the reference genome sites covered by at least 1 read, at most 4% divergence from the reference genome (computed as the fraction of non-reference alleles), a maximum of 4.5% estimated proportion of sites affected by cross-contamination, a ratio of modal coverage on the X chromosome to that on the 3RL chromosome of between 0.4 and 0.6 (male) or between 0.8 and 1.2 (female), and a genetic distance of at least 0.006 between all pairs of individuals, to exclude excessively similar samples (fig. S1a, Supplementary text). In total, 665 individuals (79%) passed the QC steps outlined above. Out of the 173 samples that were removed, 36 fell outside the Funestus Subgroup mitochondrial haplogroup (fig. S9e).

After this per-individual QC, we performed QC on the dataset as a whole, in order to check for outliers and potential sample swaps. Dataset QC was performed on all samples passing the first QC step, using principal component analysis (PCA) of the 2L chromosomal arm, which in *An. funestus* does not contain any large, common chromosomal inversions (*19*). The identification of outliers within the dataset was based on their isolation from other samples in the first 11 principal components. Identified outliers were removed and the PCA was recomputed on the remaining samples, until no outliers were identified. We excluded nine samples in two rounds of PCA, resulting in a total of 656 samples remaining for further analysis (table S1).

##### Historic samples

Historic samples have varied, but generally low coverage and hence we relaxed the QC thresholds and only applied the divergence filter; we excluded 30 individuals with > 4% divergence from the reference genome and retained 45 (table S4). As a dataset QC substitute, we performed a PCA with historic and modern samples (fig. S9c); all individuals fell within the expected clusters.

##### Public Datasets

For analyses including publicly available samples from the Funestus Subgroup (*21*) and the Folonzo and Kiribina ecotypes (*23*), we aligned all samples from these studies to the AfunGA1 reference genome and performed variant calling and quality control as described above. We compared gene drive target availability in *An. funestus* with 762 Gambiae Complex mosquitoes (*16*) and assessed variation in the *dsx* target in 3,081 individuals from the Gambiae Complex (*72*).

##### Site filters

Following QC, we implemented a site filtering procedure to address the inherent variation along the genome in our ability to confidently call genotypes. We computed various site statistics from the data of all females passing sample QC. We generated two distinct site filters, namely the static-cutoff (sc) and decision-tree (dt) filters (fig. S1c,d, table S2, Supplementary text).

For most of our analyses we used the sc filter, which retains sites with mean genotype quality (GQ) of at least 80, mean mapping quality (MQ) of at least 50, and median genomic coverage of samples with data at the focal site between 30x and 40x (fig. S1b). Depending on the analysis, further filtering based on the fraction of samples missing data was completed. These thresholds were chosen by considering the observed distributions of these statistics (fig. S1b).

For all haplotype based analyses, we used the dt filter because we completed haplotype phasing using this filter. The decision tree was trained on 15 lab colony crosses and approximately 2,000 wild-caught *An. coluzzii* and *An. gambiae* mosquitoes (*73*). Mendelian errors were computed for the colony crosses and split into training and validation data. The decision tree inputs were 15 site summary statistics computed on the wild-caught mosquitoes and the optimal tree was selected using the validation data. Because the decision tree takes only site summary statistics as input, the trained model can be applied to a different population, a different species, or a different reference genome. We applied the trained model to the site summary statistics computed from all female samples passing QC to generate the dt site filter for *An. funestus.* It resulted in fewer variant sites than the sc filter, though overall concordance was high.

##### Variant annotation

We extracted features from the AfunGA1 reference genome, using annotations from Vectorbase gff3 version 61 (*74*). Within each canonical coding sequence, we assess the effect of all possible SNPs, classify them as ‘synonymous’ or ‘non-synonymous’ mutations and record the amino acid changes for the latter category, using the CodonTable module in biopython (*75*). The AfunGA1 annotation gff3 file has not been manually curated, and we have noted some inaccuracies (Supplementary text).

#### Haplotype phasing

We performed haplotype phasing on genomic sites that met three criteria: a) passed our dt site filtering process, b) were biallelic in our dataset, and c) contained the reference allele. To manage this large dataset, we first created interval tables for each chromosome, which defined intervals of 200,000 single nucleotide polymorphisms (SNPs), with a 40,000 SNP overlap between adjacent intervals. We also incorporated a genetic map into the phasing process, which detailed recombination rates (2.0 cM/Mb for euchromatic and 0.5 cM/Mb for heterochromatic regions) inferred from average values in *An. gambiae* (*16*). The haplotype phasing process was carried out using a specialised pipeline developed by the Broad Institute’s Data Engineering team (*76*), with computational tasks conducted on the Terra platform (*77*). We applied two phasing methods: read-backed phasing conducted with WhatsHap v1.0 (*78*) and statistical phasing carried out using SHAPEIT4 v4.2.1 (*79*), successfully phasing all 656 individuals that passed our sample and dataset QC.

#### CNV calling

In the 2RL:8.2-9.8 Mbp region surrounding the *rp1* locus, we conducted copy number variation (CNV) analysis following the method described in Lucas et al. (*80*). Briefly, for each individual, we record read counts in 300 bp non-overlapping windows and normalise by the per-individual mean number of reads in genome-wide autosomal 300 bp windows, stratified by the GC content. These normalised coverage values were used as the observations in a Gaussian Hidden Markov Model (HMM) with the copy number states as hidden variables. CNVs were defined as having at least five consecutive 300 bp windows with elevated HMM-predicted copy number states. Using unique patterns of discordant read pairs and split reads at the CNV breakpoints, we manually characterised nine CNV alleles. We say an identified CNV allele is present in an individual, if we find at least two supporting diagnostic reads.

#### Cohort definitions

For the 656 samples that passed all quality control (QC) filtering we defined two sets of cohorts: *geographic cohorts* based on collection metadata, and *PCA cohorts* based on observed structure in PCA projections (Fig. 1 legend, table S1). Geographic cohorts were defined following the same strategy used in MalariaGEN Vector Observatory (*81*), where we use the coarsest administrative subdivision within each country to assign samples to cohorts based on their collection location. We did not incorporate a time component in the cohort division, because its influence on the structure of this sampleset is vastly outweighed by the location component and all samples were collected within four years of each other. The PCA cohorts were defined based on the structure observed in the PC projection of chromosome arm 2L (Fig. 1a). Each PCA cohort contains one or multiple geographical cohorts in their entirety, with the exception of GH-N. In this case, nearly all (35/36) samples formed a separate cluster on the PCA plot and were assigned to the PCA cohort North Ghana, while one sample clustered with the Equatorial PCA cohort and was not assigned to any PCA cohort. PCA cohorts were used in analyses where the advantage of a larger sample size outweighs the disadvantage of reduced homogeneity within a cohort, e.g. analyses where cohorts are further divided by inversion karyotype.

##### Sample subsets

As a default, we perform analyses on all individuals that passed QC and refer to them as **subset_1**; if specific cohorts are excluded from subset_1, this is stated explicitly. We observed an excess of heterozygous calls in samples with median coverage below 20x and suspect this is a technical artefact (Fig. 1b, fig. S2a). Analyses that estimate genetic diversity, e.g. nucleotide diversity, are sensitive to samples with outlier heterozygosity, so these analyses were performed only on individuals with median coverage ≥20x (619 samples), referred to as **subset_2**. Some analyses (e.g. Tajima’s D) are additionally affected by cohort sizes. For those analyses, we select 30 samples from each geographic cohort with median coverage ≥20x and the lowest contamination estimates. These 390 samples are referred to as **subset_3**. Four geographic cohorts (CF, KE-W, TZ, MW), have fewer than 30 samples and are thus not represented in subset_3.

#### Population genetic and selection analysis

Most analyses were performed within the MalariaGEN computational environment using scikit-allel v1.3.5-8 (*70*) functions. Analyses were performed using the sc filter, unless stated otherwise. Maps were generated with SciTools/cartopy v0.22.0 (*82*), and other plots were generated with matplotlib (*83*).

##### PCA

PCAs were computed using the pca function on a random sample of typically 200,000 biallelic sites within the specified region, with a minor allele frequency ≥0.01 and less than 5% of samples with missing genotype calls.

The sliding window PCA (inspired by (*84*)) uses the `pca` function within windows of 5000 variants, moving with 1000 variant steps. We selected nearly-biallelic variants (defined as having second minor allele frequency ≤0.001) with a minor allele frequency ≥0.02, allowing for at most 10% of samples with missing data, and downsampled to an average of 5000 variants per Mbp. This subsetting of variants was necessary to include sufficient sites along the genome. Continuity between windows is ensured by minimising the absolute distance of PC1 values between the focal and preceding window, flipping the PC1 axis if required.

For PCAs including historic samples, we excluded C>T and G>A substitutions from our variant selection to avoid confounding signals from DNA deamination. This correction was not implemented for the sliding window PCA, because it would substantially increase the computation time.

##### Nucleotide diversity and Tajima’s D

We estimated nucleotide diversity (π) as the mean pairwise differences for subset_2 individuals in each geographic cohort using the `mean_pairwise_difference` function. We split the genome into non-overlapping windows of 20,000 accessible sites with genotype calls for ≥90% of samples in each cohort and averaged the values within each window. Tajima’s D is affected by sample size and smaller cohorts tend to have higher (less negative) Tajima’s D values (data not shown). We partitioned the genome into windows as above, but using individuals from subset_3, ran the `tajima_D` function and averaged the values within each window.

##### F_ST_

We followed Hudson’s method (*85*) to estimate pairwise F_ST_ values. We computed F_ST_ using the `hudson_fst` function, which gives the numerator (number of differences between cohorts minus number of differences within cohorts) and denominator (number of differences between cohorts) for each site (for non-variable sites both are equal to 0 and hence the F_ST_ value for these sites is undefined). As per recommendation (*86*), we reported the “ratio of averages” per chromosomal arm (mean of numerators divided by the mean of denominators).

##### Heterozygous sites and runs of homozygosity (ROH)

We used the count_het function to count the number of heterozygous sites for every individual. We computed ROHs only for females (21 males excluded) on genomic sites that passed the dt filter using the `roh_mhmm` function with `min_roh=100000` and default parameters otherwise. This function utilises a multimodal hidden Markov model (MHMM) to estimate the positions and lengths of ROHs.

##### Doubletons

We identified doubletons as alleles at accessible sites occurring exactly twice in subset_3 using the count_alleles function. We then identified the number of doubletons shared within and between all geographic cohorts. Some of these doubletons will be caused by recurrent mutations and thus not have shared ancestry.

##### In silico inversion karyotyping

Initial karyotyping was performed by identifying two horizontal thresholds in the sliding window PCA, partitioning the samples into three groups. To account for the effect of geographic structure, we used two different sets of thresholds. We determined the standard orientation by incorporating AfunGA1 as a fully homozygous sample, representing the 2R+^t^+^a^h, 3Rab, 3La orientation. We checked the results by computing PCA on the entire inversion region and assess whether samples clustered by karyotype; by this procedure we assigned karyotypes for GH-N, which is not included in the sliding window PCA, for all inversions except 3La.

##### Mitochondrial tree

We extracted reads mapping to scaffold_MT of AfunGA1 into separate BAM files, and used bcftools 1.10.2 (*66*) `mpileup` to create consensus mitochondrial sequences for each sample. Together with publicly available Funestus Subgroup and sister species mitochondrial sequences(*20*, *21*) (table S1), we created a maximum likelihood tree using MAFFT v7.520 (*87*) and FastTree v2.1.11 (*88*), and visualised it using TreeViewer v2.2.0 (*89*).

##### Garud’s H_12_ scan, haplotype trees, and variants potentially under selection

We conducted Garud’s H12 scans (*35*) using the `moving_garud_h` function with cohort-specific window sizes (table S2). H12 was performed on sites passing the dt-filter. H12 scans were not presented for GH-N due to a high noise-to-signal ratio caused by a high number of ROHs. For regions with H12 ≥ 0.4, we defined a peak as a 0.2 Mbp region centred on the highest value. Next, we calculated SNP allele frequencies for all non-synonymous variants (without site filter) for each gene in this region with minor allele frequency ≥ 0.055 using the `snp_allele_frequencies` function (table S3). Within the peaks, we identified genes or gene families likely to be under selection and constructed haplotype trees using the `plot_haplotype_clustering` function.

For historic samples, the genotype at potential insecticide resistance mutations was confirmed in IGV v.2.17.1 (*90*), as DNA deamination can be erroneously called as variants, especially at low coverage.

##### Gene-drive targets

We identified gene-drive targets in the reference genome as 20 bp sequences entirely within coding sequence, (using Vectorbase gff version 65 (*74*)) containing the protospacer adjacent motif (ending in `-GG` on the + strand or starting with `CC-` on the - strand). To take into account resistance due to natural variation at the target site, we eliminated all gene-drive targets that contained any variants in the group of samples under consideration (from subset_2).

##### Doublesex

The doublesex (*dsx*) target site is found at 2R:48,714,637-659 in AgamP4 and at 2RL:15,613,532-554 in AfunGA1. We searched for any variants at these sites in 3081 Gambiae Complex individuals (*72*) and *An. funestus* subset_1 individuals.

### Supplementary Figures

**Fig. S1.**
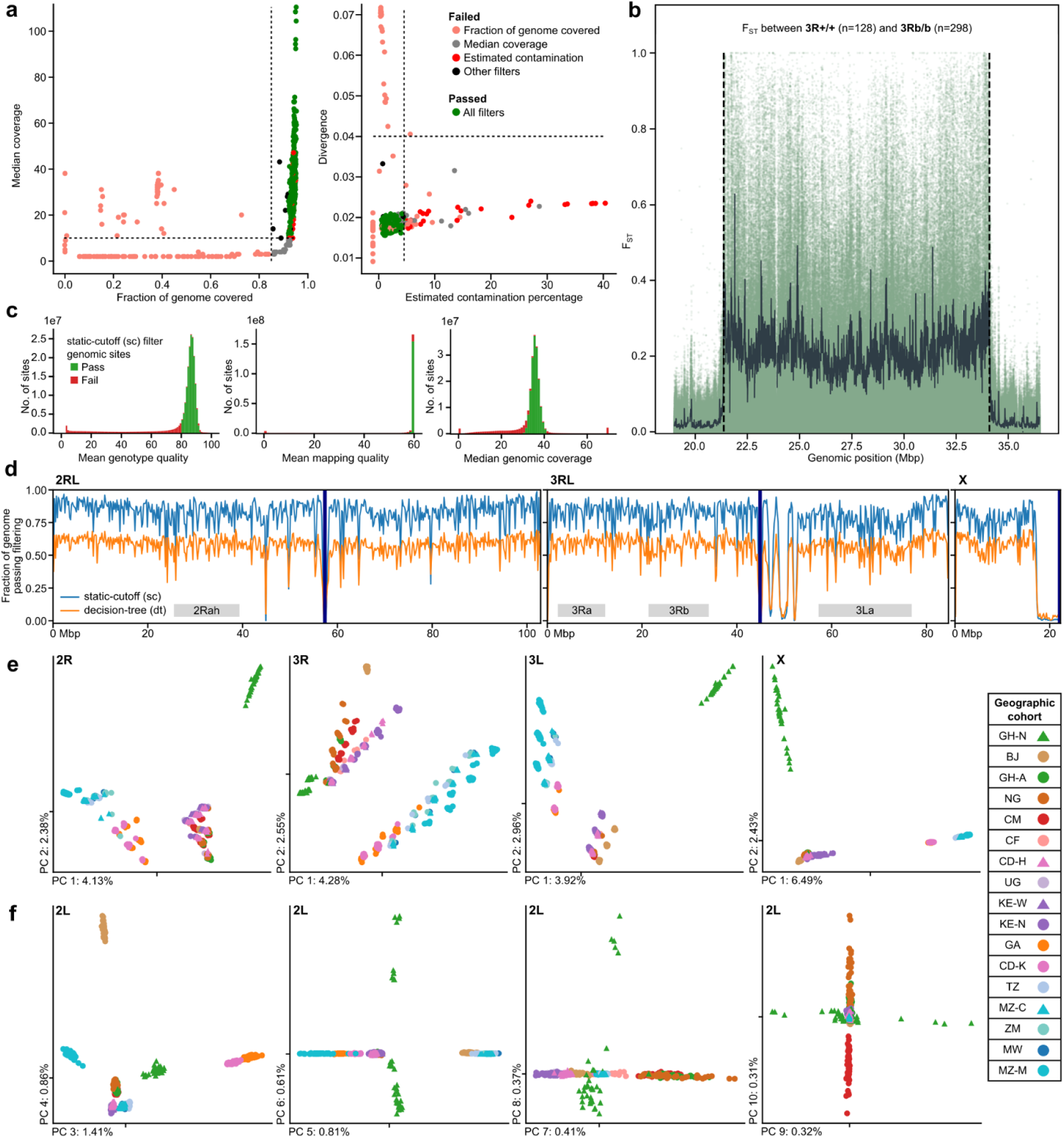
Quality control assessment of *An. funestus* individuals in this study. (**a**) Values of four quality control statistics for 838 sequenced, aligned, and variant called samples. Samples are coloured by iteratively applying stated thresholds in the order they appear in the legend and the colour represented is the final status of the sample (see Methods for details on excluding samples during the QC process). (**b**) F_ST_ between homokaryotypes for the 3Rb inversion was used to infer approximate inversion breakpoints; light green dots are values for single variants, dark green line is the 6 kbp sliding window average, dashed vertical lines are the inferred breakpoints. (**c**) Site summary statistics used for the static-cutoff (sc) filter, computed on females passing QC (636 individuals). We required sites to have mean genotype quality (GQ) ≥80, mean mapping quality (MQ) ≥50, and median coverage (excluding samples with 0 coverage at the particular site) between 30x and 40x. Sites passing these criteria are shown in green, sites failing in red. (**d**) Fraction of accessible sites in 200 kbp windows for the sc and decision-tree (dt) site filters. Centromeres are indicated by dark blue vertical lines and inversions as horizontal grey boxes. (**e**) Projection of individuals coloured by geographic cohort along the first two principal components (PCs) on chromosome arms 2R, 3R, 3L and X. Percentage of explained variance is shown in the axes labels. (**f**) Projection of individuals coloured by geographic cohort along PCs 3 to 10 on chromosome arm 2L. Percentage of explained variance is given in the axis labels.

**Fig. S2.**
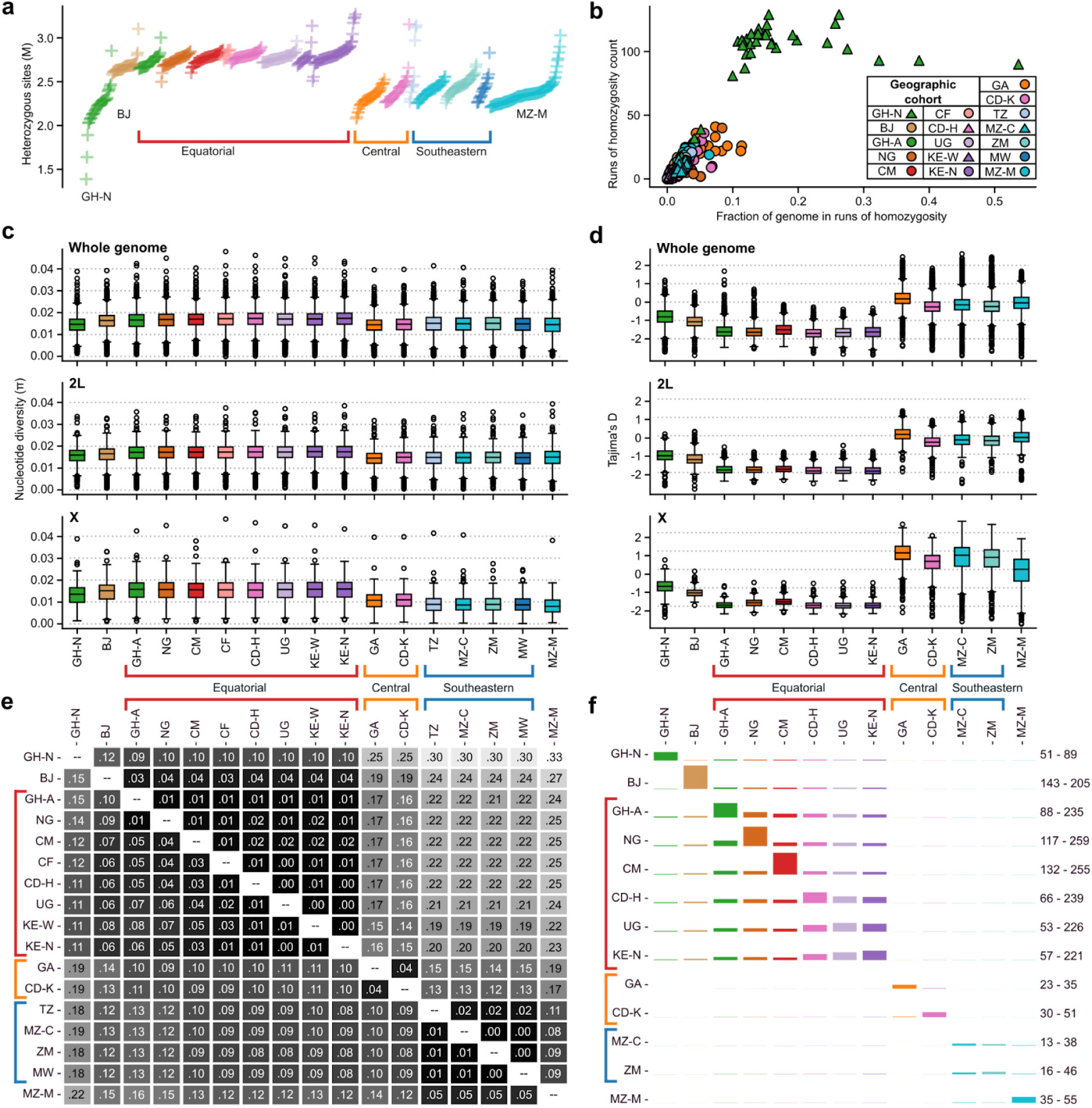
Genetic diversity in 17 *An. funestus* cohorts from this study. (**a**) Number of heterozygous sites per individual in subset_1 (females only) on all accessible sites coloured by geographic cohort. Individuals are displayed as crosses and ordered along the horizontal axis by geographic cohort and increasing number of heterozygous sites along the vertical axis (number in millions of sites). PCA cohorts containing multiple geographic cohorts are indicated by square brackets for this and subsequent subplots. (**b**) Runs of homozygosity (ROH) for female mosquitoes from subset_1 (21 males excluded) where the fraction of the genome contained in ROHs of length ≥100 kbp is plotted on the horizontal axis against the total count of ROHs on the vertical axis. (**c**) Box plots of nucleotide diversity (π) per geographic cohort (using individuals from subset_2), computed in non-overlapping windows of 20 kbp accessible sites. Aggregated over all nuclear accessible sites (top), sites on chromosome arm 2L (middle) and on chromosome arm X (bottom). (**d**) Box plots of Tajima’s D per geographic cohort (using individuals from subset_3, which excludes cohorts CF, KE-W, TZ and MW with fewer than 30 individuals), computed in non-overlapping windows of 20 kbp accessible sites and aggregated over all accessible sites (top), accessible sites on 2L (middle) and on X (bottom). (**e**) Pairwise fixation indices (F_ST_) between pairs of geographic cohorts (subset_2). Computed on all accessible sites on 2L (lower triangle) and X (upper triangle). (**f**) Doubleton sharing patterns between geographic cohorts (subset_3). Each row shows the distribution of the other allele of a doubleton per geographic cohort, given that at least one allele of the doubleton is found in the focal cohort represented by that row (so effectively each doubleton is counted twice, once for each individual in which the doubleton is found). The numbers on the right indicate the amount of doubletons shared within the focal cohort (×10,000) and the total number of doubletons with at least one allele found in the focal cohort (×10,000), respectively.

**Fig. S3.**
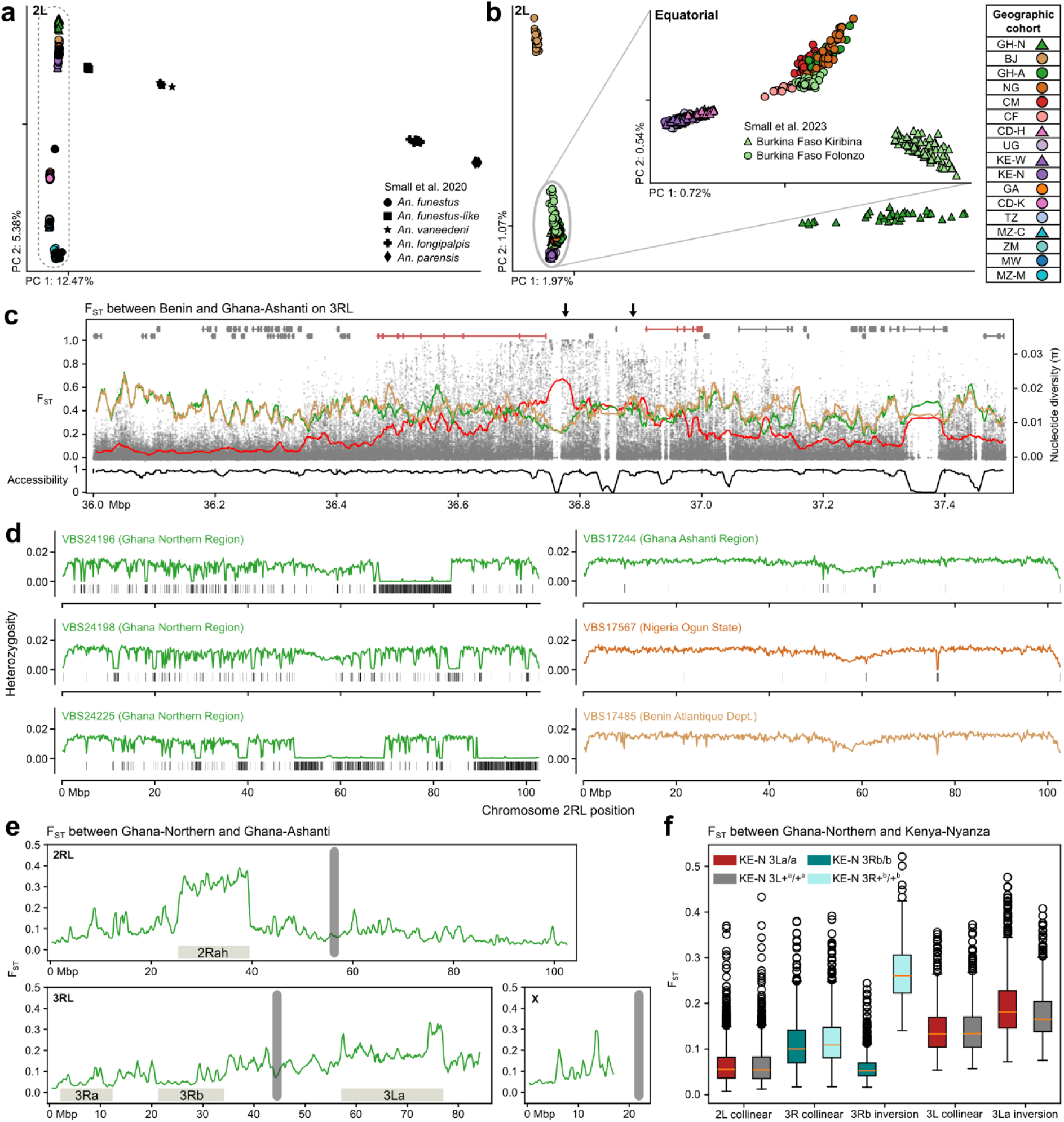
Ecotypes and differentiated populations. (**a**) Principal component projection of chromosome arm 2L containing all our samples as well as other species from the *An. funestus* subgroup (*21*). All samples from this study fall within the dashed line area, containing only *An. funestus* s.s. (**b**) Principal component projection on chromosome arm 2L computed on samples from the Equatorial, South Benin and North Ghana PCA cohorts, as well as Folonzo and Kiribina ecotypes from Burkina Faso (*23*). The inset is computed only on the samples from the Equatorial cohort and those from Burkina Faso. (**c**) F_ST_ between Benin Atlantique Dept (BJ) and neighbouring cohort Ghana Ashanti Region (GH-A) centred on the 3R peak of differentiation of BJ from other cohorts (Fig. 1d). Each grey dot shows the F_ST_ value of a single accessible site and the red line shows the sliding window average F_ST_ of 10 kbp accessible sites (step size 1 kbp). In ochre and green is the windowed nucleotide diversity for the two cohorts, using the same window sizes. The black line below shows the fraction of accessible sites in 10 kbp windows. On top are genes present in this region, with exons shown as vertical stripes and introns of the same gene as horizontal lines; + strand on top, - strand below. In pink are two semaphorin-2A-like protein coding genes. The arrows link to zoomed-in IGV views in fig. S10. (**d**) Heterozygosity across the 2RL chromosome in three GH-N individuals (left), and three individuals from neighbouring populations (GH-A, NG, BJ). Vertical black bars at the bottom denote detected ROHs. (**e**) F_ST_ between Ghana Northern Region (GH-N) and Ghana Ashanti Region (GH-A) in sliding windows of 500 kbp accessible sites along the genome, moving with 100 kbp steps. Inversion regions are indicated by horizontal grey bars, centromeres by vertical grey bars, as in Fig 1d. (**f**) Bar plots of F_ST_ values in non-overlapping windows of 20 kbp accessible sites between Ghana Northern Region and Kenya Nyanza Province (KE-N) on the 2L chromosome arm, the collinear part of the 3R chromosome arm, the 3Rb inversion region, the collinear part of the 3L chromosome arm and the 3La inversion region. On 2L and 3L, we compare GH-N against 3La/a and 3L+/+ homozygotes from KE-N, while on 3R we compare GH-N against 3Rb/b and 3R+/+ homozygotes from KE-N. For the collinear parts of the chromosome arms, the F_ST_ values compared to the different homozygous orientations are the same, while for the 3Rb inversion region the F_ST_ values to 3Rb/b individuals are lower than on the collinear genome, while the F_ST_ values to 3R+/+ are higher than on the collinear genome. For the 3La inversion, we do not observe such a pattern.

**Fig. S4.**
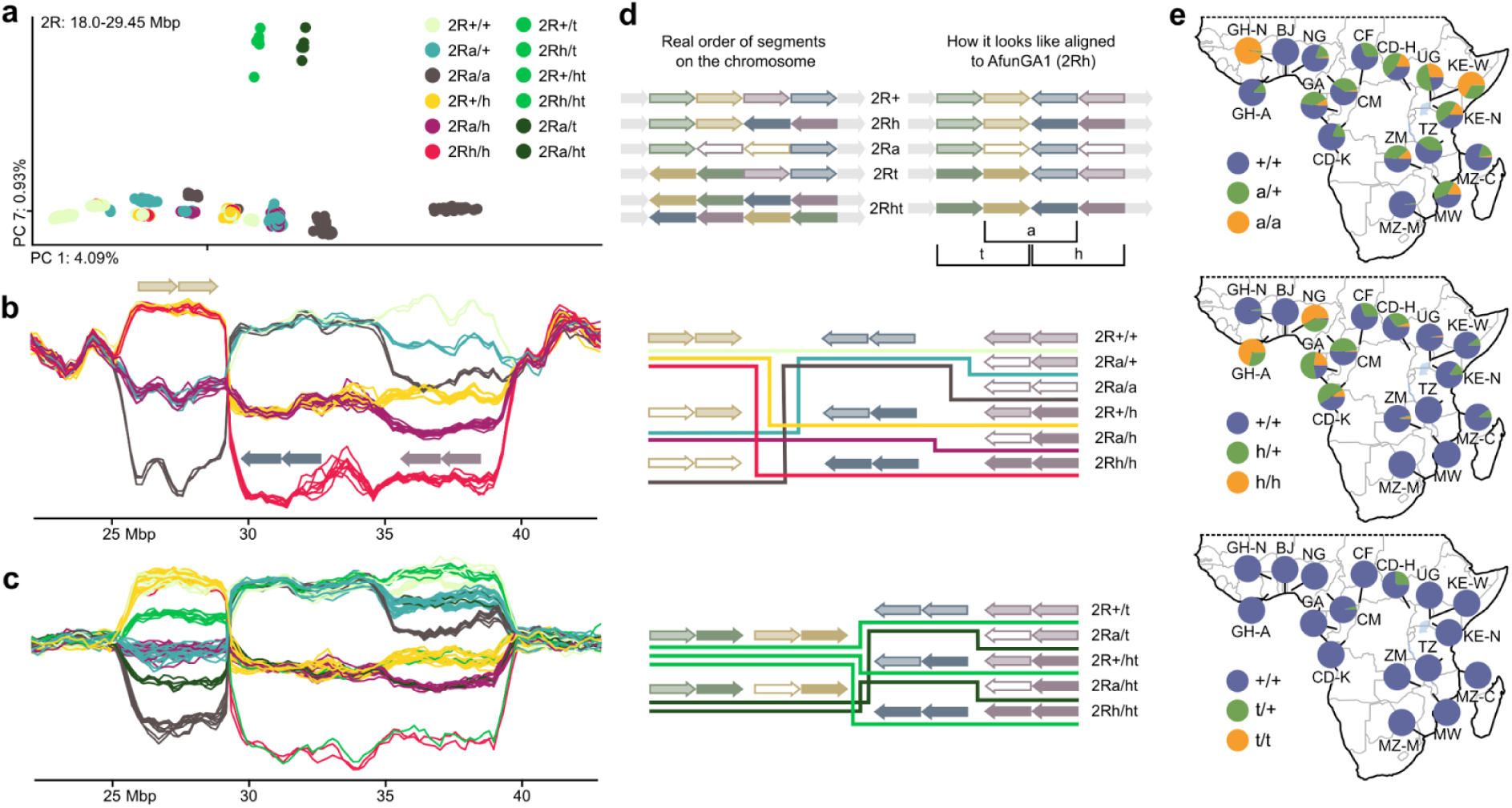
Overlapping inversions (a, h, t) on the 2R chromosome arm. (**a**) Projection along the first and seventh principle components on the 2Rt inversion region computed on 100,000 randomly selected variants. Samples are coloured by their inferred combined karyotype for the 2Rt, 2Ra and 2Rh inversions. (**b**) Sliding window PC1 on the region of overlapping inversions 2Ra and 2Rh, here shown for Gabon (GA) samples only. The 2Ra inversion region is broken up into two pieces when aligned to AfunGA1. Arrows denote genomic regions in a sample as mapped to the AfunGA1 reference (see panel d). There are six possible combined karyotypes for 2Rah; here the samples are coloured by their inferred combined karyotype as in panel a. (**c**) Sliding window PC1 on the region of overlapping inversions 2Ra and 2Rh, here shown for Cameroon Adamawa (CM) and DRC Haut-Uélé (CD-H). On the part of the 2Ra inversion region where it is not overlapped by 2Rh (∼Mb 26-29) there are two additional karyotype bands (in light and dark green) falling in between the three bands also observed in panel b. The samples in these bands are heterozygous for a third inversion on the 2R chromosome arm, 2Rt. Only 13 samples in the entire dataset are heterozygous for 2Rt: two from Cameroon and 11 from DRC Haut-Uélé, and there are no samples homozygous 2Rt/t. There are 11 observed combined karyotypes for 2Raht, where 2Rt only occurs as homozygous standard or heterozygous. (**d**) Schematic overview of the observed overlapping inversions on 2R. Top: believed order of DNA segments within the 2R inversion on the physical chromosome and alignment of the orientations to the AfunGA1 reference genome. There are two ways of ordering the segments for 2Rht on the physical chromosome, but they both result in the same alignment. Middle: ignoring the 2Rt inversion, there are three distinct chromosomal orientations resulting in six possible karyotypes. Each karyotype corresponds to one of the trajectories observed in the sliding window PCA. Bottom: 2Rt is only observed in its heterozygous state. Restricting our attention to the 2Rt heterozygous individuals, there are two orientations containing 2Rt and three orientations without 2Rt, so there are six possible karyotypes where 2Rt is in a heterozygous state. However, it is impossible to distinguish between 2R+/ht and 2Rh/t using only genotype information, as the sliding window PC1 trajectories are the same. So we observe five additional trajectories in the sliding window PCA for samples that are heterozygous for 2Rt. (**e**) Distribution of 2R inversion karyotypes for each geographic cohort.

**Fig. S5.**
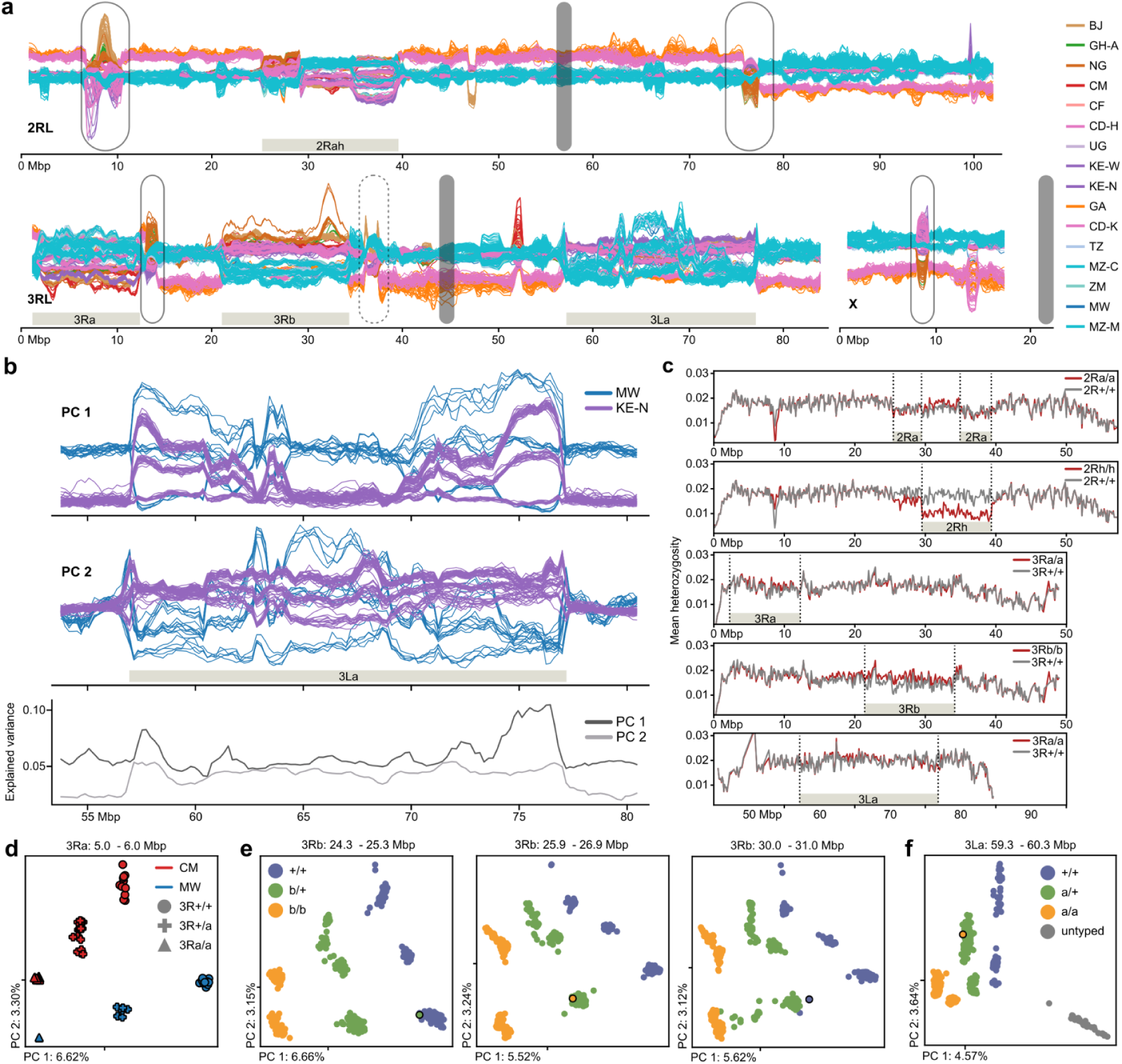
Recombination and heterozygosity inside inversion regions. (**a**) Sliding window PCA displaying PC2 along the genome for all individuals except GH-N (complementary to Fig. 1d). Plot annotations as in Fig. 1d. (**b**) Sliding window PCA in the 3La inversion region computed on all cohorts except GH-N, but displaying only MW and KE-N for visibility. The top panel shows the PC1 values, the middle panel the PC2 values, and the third panel the fraction of variance explained by PC1 and PC2 in dark and light grey respectively. (**c**) Mean heterozygosity of homokaryotypes from the Equatorial PCA cohort in 100 kbp non-overlapping windows for five inversions (2Ra, 2Rh, 3Ra, 3Rb, 3La). The red line is homozygous inverted, the grey line is homozygous standard. The region of the focal inversion is indicated by horizontal grey bars and the breakpoints by dashed vertical lines. (**d**) Projection along the first two principal components computed on all samples on a 1Mb region within the 3Ra inversion. For visibility, only CM and MW are shown, as in Fig. 2a. Colours indicate geographic cohort, shapes indicate inversion karyotype. Note that CM 3R+/+ and MW 3R+/a overlap on PC1 but are differentiated on PC2, due to the compound signal of geographic structure and inversion karyotype. (**e**) Projection along the first two principal components computed on all samples on 1 Mb regions within the 3Rb inversion. Regions are centred on the peaks of the putative double recombinants (bold lines changing trajectories in Fig. 2b); in this figure those samples are highlighted with a black outline. (**f**) As in (e) but for a double recombinant in the 3La inversion.

**Fig. S6.**
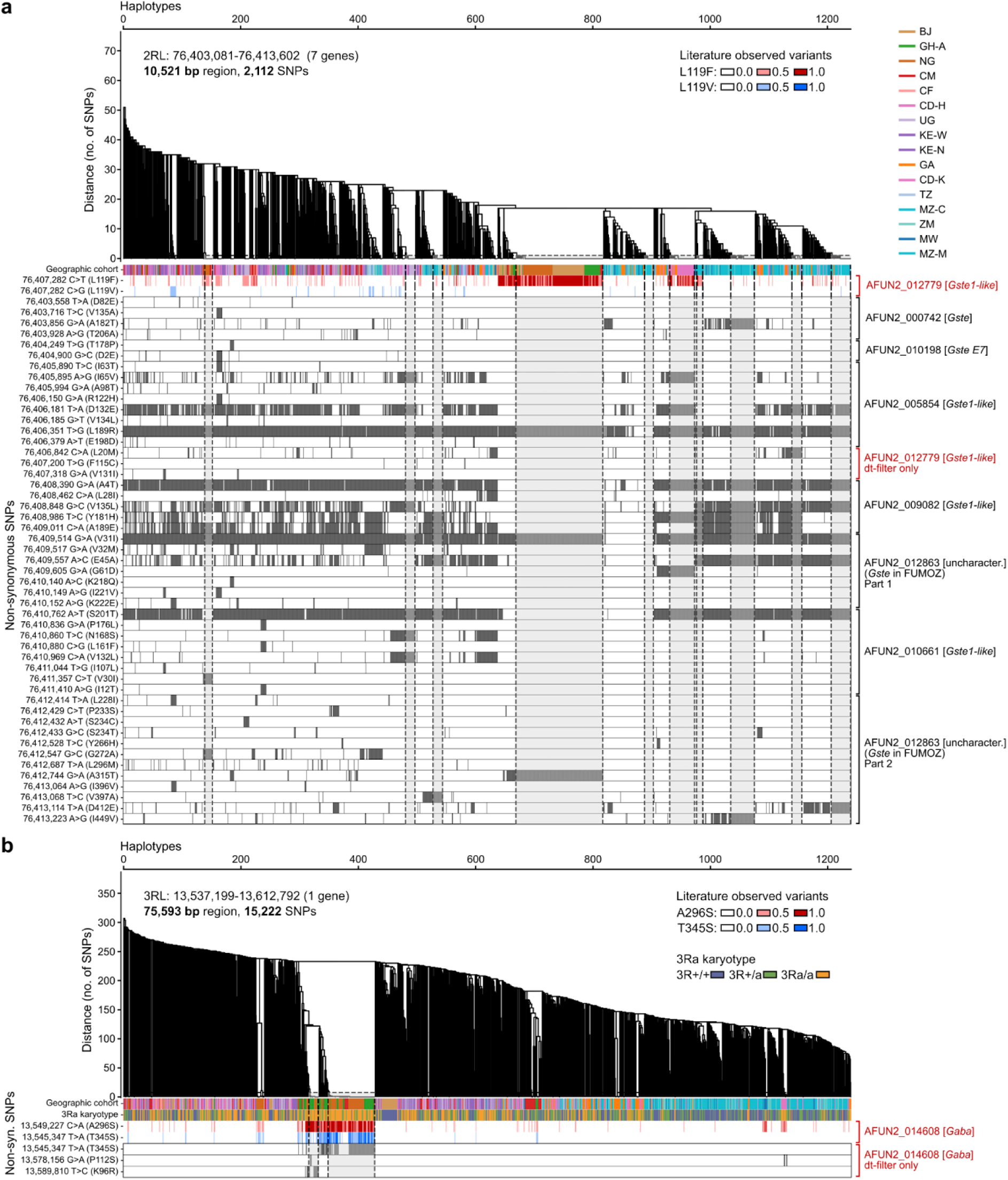
Full haplotype clustering trees for *Gste2* and *Gaba*. (**a**) Haplotype clustering within a region containing seven GSTe genes. The dendrogram is obtained by hierarchical clustering of phased haplotypes, and used to define haplotype clusters as groups of haplotypes with SNP divergence below 0.0005 (cutoff indicated as dashed horizontal line on the dendrogram). The first bar below the dendrogram shows the population of origin for each haplotype. Next, the red bar shows the known GSTe2 L119F mutation, and the blue bar the previously unreported L119V mutation (note that the sites of these mutations were filtered out before haplotype phasing, so each haplotype is coloured by the genotype of the individual it belongs to). The bars below show the presence (grey) or absence (white) of each non-synonymous mutation with maf ≥ 0.055 that was included in haplotype phasing. The genes in which these mutations occur are listed on the right (known resistance gene in red, highlighted in Fig. 3). Haplotype clusters are indicated by grey shaded areas within dashed grey lines. (**b**) Haplotype clustering within the *Gaba* gene, same structure as panel a, with the addition of a horizontal bar depicting the 3Ra karyotype present upstream from the gene (coloured by the karyotype of the individual), and the two known *Gaba* non-synonymous variants A296S (red) and T345S (blue).

**Fig. S7.**
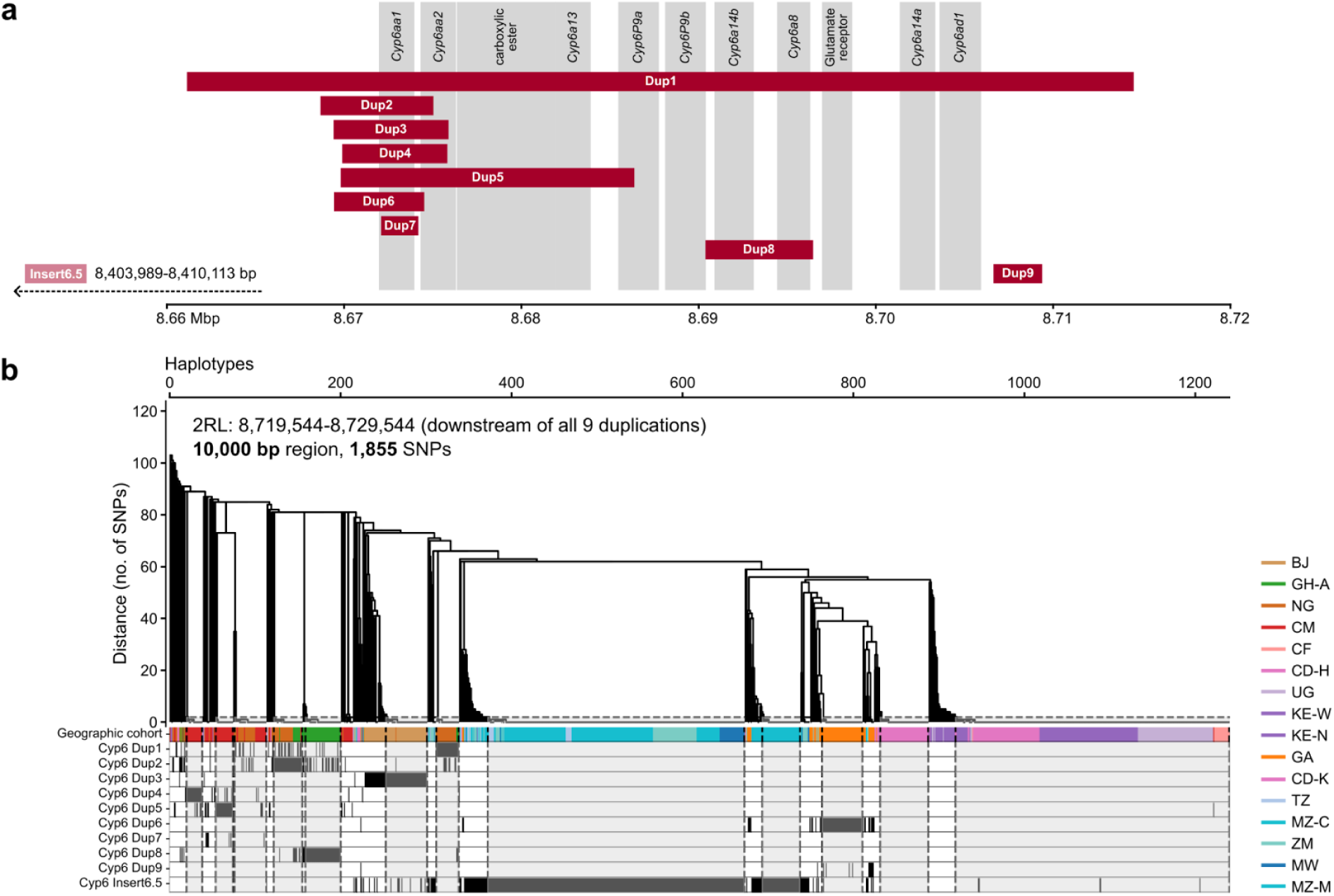
Haplotype clustering tree for *Cyp6p* (Rp1) with copy number variants. (**a**) Approximate locations of detected duplications on the 2R arm within a cluster of cytochrome P450 genes. The red bar indicates the genomic region that is duplicated in relation to the reference genome. (**b**) Haplotype clustering within a 10,000 bp region downstream of the cluster of Cyp6 genes and of the identified duplications (haplotype clustering on the region containing the Cyp6 genes was affected by the CNVs, see Supplementary text). The dendrogram is obtained by hierarchical clustering of phased haplotypes based on SNP distances and the first bar shows the population of origin for each haplotype. The bars below show the presence (black) of observed copy number variants (CNVs) relating to specific duplications upstream of the region used for haplotype clustering. CNVs are called as present or absent for each individual, so each haplotype is coloured by the CNV status of the individual it belongs to. Haplotype clusters, where all haplotypes share exactly the same SNPs, are indicated by grey shaded areas within dashed grey lines.

**Fig. S8.**
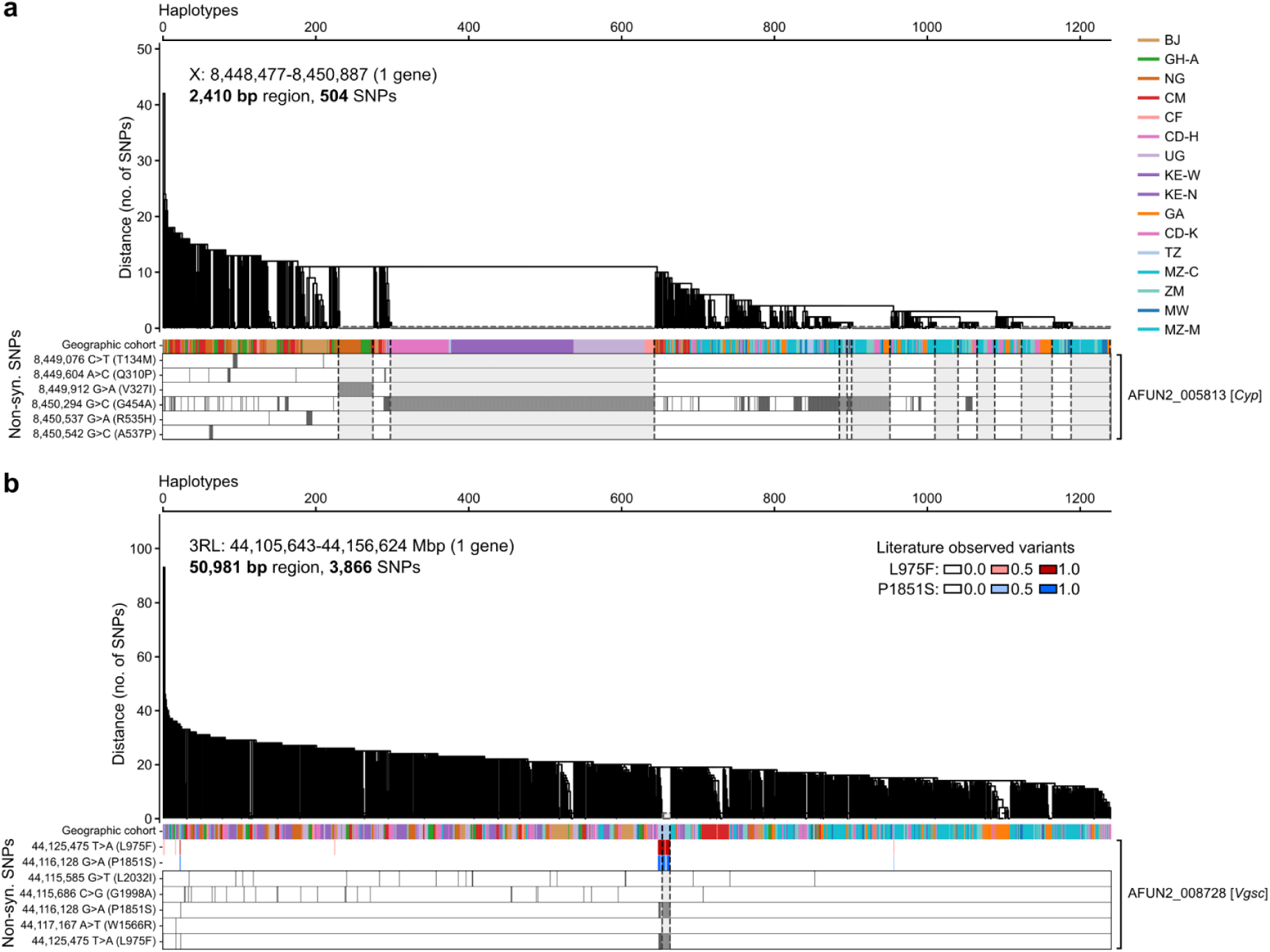
Haplotype clustering trees for *Cyp9k1* and *Vgsc* genes. (**a**) Haplotype clustering within the *Cyp9k1* gene, same depiction as fig. S6. (**b**) Haplotype clustering within the voltage gated sodium channel (*Vgsc*) gene, same depiction as fig. S6. The red bar shows the known *kdr* L975F mutation, and the blue the P1851S mutation, which is present at a similar frequency as L975F, but is not a known resistance mutation.

**Fig. S9.**
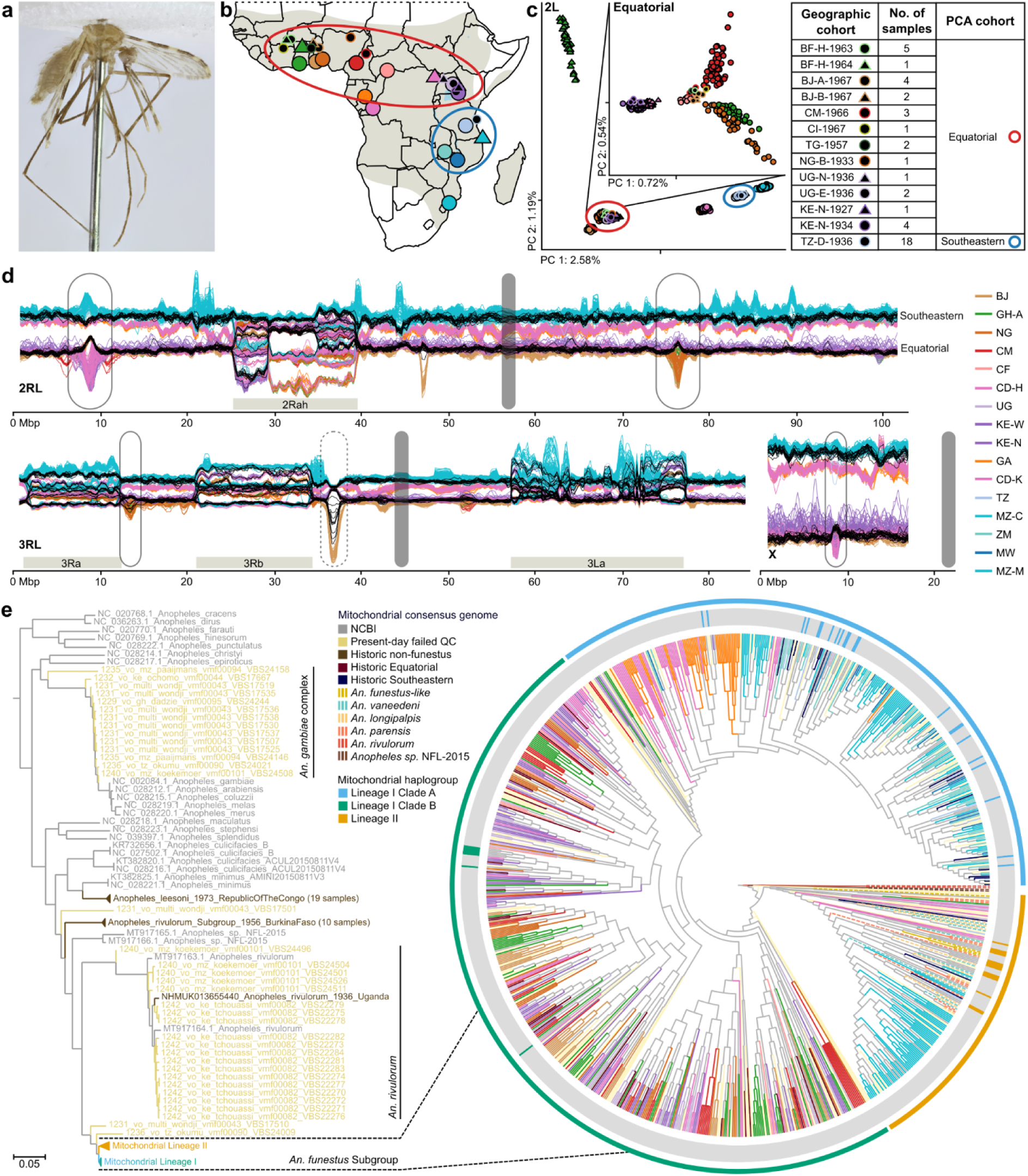
Exploration of historic *An. funestus* specimens. (**a**) Canon 5DSR focus stacked image of historic pinned specimen NHMUK014063606 (collected by Major H. S. Leeson 28/08/1936 in Dar es Salaam, Tanzania) after minimally morphologically destructive DNA extraction and critical point drying, returned to the London Natural History Museum collection. (**b**) Original collection location of 45 sequenced historic *An. funestus* individuals depicted by black circles superimposed over the present-day sample set. Geographic cohort names also include collection years (from 1927 to 1967). (**c**) PCA of present-day and historic (in black) individuals, showcasing that historic samples fall into the present-day Equatorial and Southeastern PCA cohorts. The inset showcases only the Equatorial cohorts, with historic individuals with a median genomic coverage <15x removed as they added too much noise. (**d**) Sliding window PC1 of the full dataset consisting of historic (black lines) and present-day *An. funestus* individuals (one historic sample with coverage <5x removed). Plot annotation as in Fig. 1d. Interestingly, a few historic samples follow the peak specific to South Benin (see fig. S10). (**e**) Consensus mitochondrial tree with 171 NCBI available Anopheline mitochondrial genomes (table S1) and all 838 present-day and 75 historic specimens (before QC). On the left is a sub-branch showcasing samples that turned out to be different species (among others Gambiae Complex and *An. rivulorum*). On the right is a cladogram zooming in on the Funestus Subgroup only (with *An. rivulorum* as outgroup). The middle circle colours NCBI individuals by mitogenome lineages as defined in Jones *et al.* (*20*), and the outer circle shows the borders of each mitogenome lineage across the dataset.

**Fig. S10.**
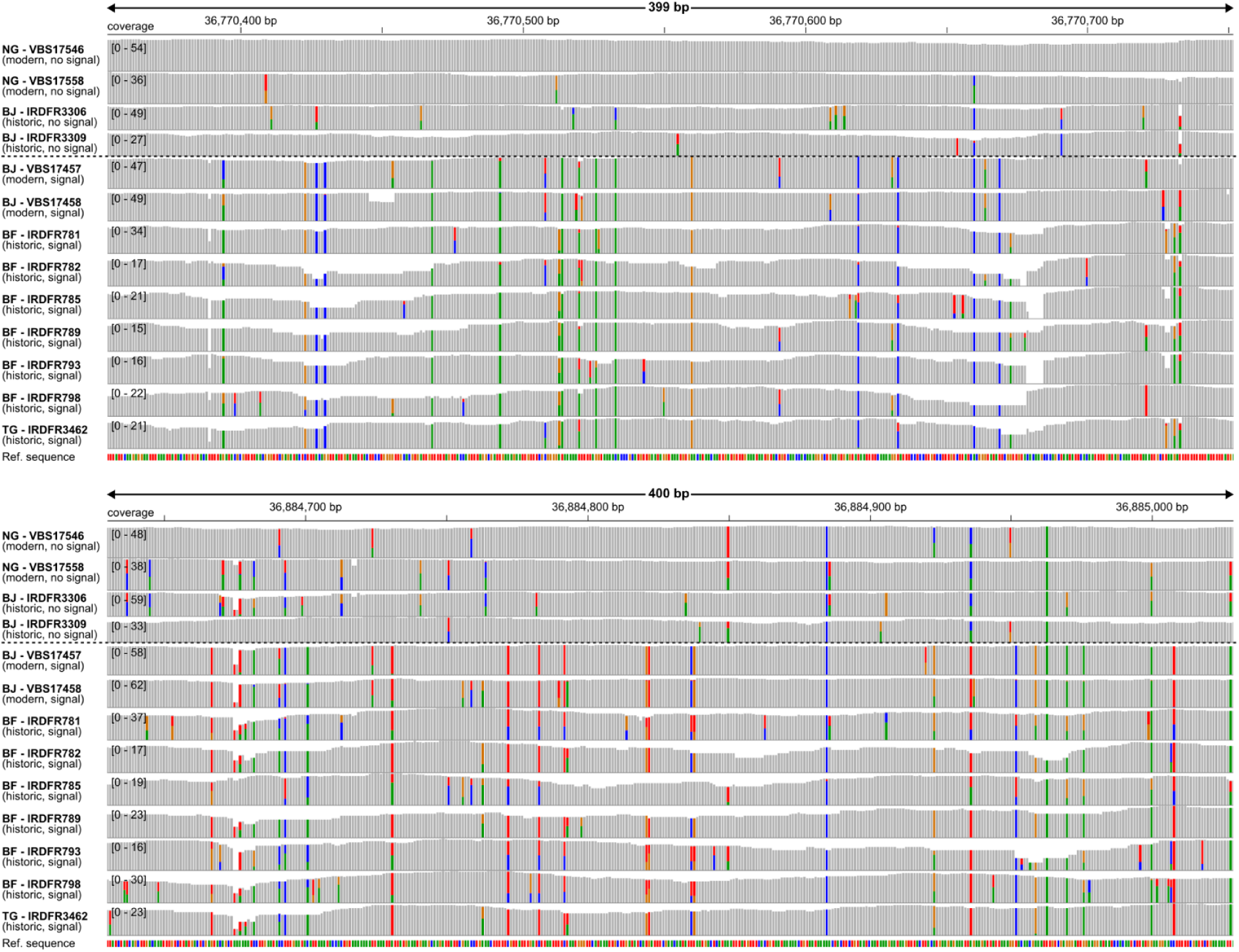
IGV views of the peak of differentiation of the South Benin cohort on chromosome arm 3R. Panels show two representative regions of 400 bp within the differentiation peak (genomic locations indicated by arrows in fig. S3c). The samples above the dotted line in each panel are modern and historic samples that do not display the differentiation signal in the sliding window PCA, while those below the dotted line do (Figs. 1d, 3c, fig. S9d). Both panels show a number of variants that are fixed between these two groups of samples.

### Supplementary Text

#### Quality Control Metrics

The exact definitions of the metrics used for QC are detailed below. Median coverage determination involved calculating depth of coverage at each genomic position, and samples with median coverage below 10x across the entirety of the genome were excluded from further analysis (ranging between 3x and 108x). The fraction of genome covered was evaluated by considering the ratio of sites with at least 1x coverage to the total genome length, and samples with a value below 85% were excluded (ranging from 0.3% to 95%). Divergence from the reference genome was determined by computing the sum of non-reference alleles divided by twice the total number of alleles called across the entire genome and samples with values greater than 0.04 were considered to be significantly differentiated from our reference genome and excluded. Contamination between samples was estimated using a previously described method (*91*), and involves computing the likelihood of observed allele counts for varying levels of contamination, where a maximum likelihood (ML) value represents the estimated proportion of sites affected by cross-contamination, and we excluded samples with ML values exceeding 4.5% due to contamination concerns. Replication likelihood was assessed to identify samples that were inadvertently sequenced multiple times (e.g. from different body parts), and was achieved by computing pairwise genetic distances between all sample pairs for autosomal genome sites where both samples possess complete genotypes, using the city block distance metric. Samples with genetic distances less than 0.006 are considered excessively similar and would be excluded to ensure the inclusion of genetically distinct individuals, however all of our samples had values above that and thus passed the replication likelihood filter.

#### Site Filters

##### Summary statistics for the dt site filter

The following statistics were computed on all female mosquitoes passing QC and used as input to the site filter decision tree to yield the dt_20200416 site filter.

- `GQ10` number of samples with Genotype quality (GQ) 10 or lower
- `GQ30` number of samples with GQ 30 or lower
- `GQ_std` standard deviation of non missing GQ values
- `GQ_mean` mean of non missing GQ values
- `MQ10` number of samples with Mapping quality (MQ) 10 or lower
- `MQ30` number of samples with MQ 30 or lower
- `MQ_std` standard deviation of non missing MQ values
- `MQ_mean` mean of non missing MQ values
- `allele_consistency` number of samples with at least one read not consistent with genotype call
- `hi_gc_normed_cov` number of samples with high coverage, where site was >2x modal coverage for the sample, within GC bins
- `lo_gc_normed_cov` number of samples with low coverage, where site was <0.5x modal coverage for the sample, within GC bins
- `hi_cov` number of samples with high coverage, where site was >2x modal coverage for the sample, not accounting for GC bias
- `lo_cov` number of samples with low coverage, where site was <0.5x modal coverage for the sample, not accounting for GC bias
- `no_cov` number of samples with no reads present at this site, i.e., coverage 0
- `repeat_dust` sites masked by dustmasker in blast v.2.12.0 (*92*)
- `repeat_repeatmasker` sites masked by RepeatMasker v.4.1.2-p1 (*93*) with master Dfam database v.3.5 (*94*) (2021-10-08) for Arthropoda and custom repeat database built with RepeatModeler 2.0.2 (*95*) rmblast 2.11.0+ (relying on TRF 4.09 (*96*), RECON (*97*), RepeatScout 1.0.6 (*98*), RepeatMasker 4.1.2 (*93*)) - as implemented in EarlGrey v2.1 (*99*).
- `ref_n` gap sites in reference genome

##### Comparison of accessibility between site filters

The sc and dt filters classify 77.41% and 54.71% of genomic sites as accessible, and within the coding sequence these percentages are 88.98% and 67.48%, respectively (table S2, Tab ”Site Filters”). In comparison, for *An. gambiae* and *An. coluzzii* from the Ag1000G Phase 1 dataset (*16*), the dt filter classifies 72.31% of all sites and 87.89% of coding sites as accessible. For *An. funestus*, the dt filter is more stringent than the sc filter, but concordance is high with only 1.98% of sites accessible according to the dt filter, but not according to the sc filter.

#### Mitochondrial structure

We compared consensus mitochondrial genomes of all our samples with previously published mitogenomes (*20*, *21*). Of the 838 samples sequenced (including those failing QC), 36 fall outside the Funestus Subgroup (all failed sample QC on the divergence filter; most fall within the Gambiae Complex or Rivulorum Subgroup clades); the remaining samples sit within the Funestus Subgroup clade. Previous mitogenomic research on *An. funestus* classified mitochondrial genomes into three distinct haplogroups: Lineage I - Cluster A, Lineage I - Cluster B, and Lineage II (*20*). The latter is the most diverse mitochondrial haplogroup that also contains additional species from the Funestus Subgroup (*20*). When comparing consensus mitochondrial genomes from each of our samples to previously published mitogenomes (*20*, *21*), structure generally follows our PCA cohorts (fig. S9e). Central cohort individuals fall in Lineage I - Cluster A, the vast majority of Equatorial cohort individuals fall in Lineage I - Cluster B, with only a few falling at the base of Clade A (*20*). Southeastern and South Mozambique cohort individuals contain both Lineage I - Cluster A and Lineage II haplogroups. Interestingly, this deep mitochondrial divergence within the cohorts spread over Lineage I and Lineage II is not evident in the structure of the nuclear genome. This pattern of mitochondrial haplogroups across the cohorts sampled here might suggest that the mitogenomes within the subgroup originated in the south or southeast, as evident by the majority of other species in the subgroup being present primarily across the east side of the continent (*21*), and then spread and differentiated across the Equatorial region, with a second spread across central and southeastern Africa.

#### Population structure

Most population structure analysis uses a subset of all accessible sites, often using biallelic sites with a minor allele frequency above a certain threshold. In each section, we motivate and describe which sites and thresholds we chose (Methods).

##### PCAs

Projections along the first two PCs for other chromosomal arms are consistent with the clustering observed on chromosome arm 2L, but the autosomes show additional structure caused by segregating inversions (fig. S1e). These large segregating inversions are also clearly visible in a sliding window PCA (values for PC 1 and PC 2 plotted in Fig. 1d and fig. S5a respectively, more on inversions below). Additionally, the sliding window PCA reveals some genomic regions where one or several cohorts deviate from their horizontal trajectory to form a peak (indicated by black-bordered boxes); some of these peaks correspond to selective sweeps and putative ecotype differentiation (Fig. 3a). PCs 3 to 10 show different aspects of the structure in this dataset, e.g. PC 3 captures the variation that North Ghana shares with the Central cohort, PC 5 captures the variation which differentiates South Benin from the Equatorial cohort and PC 7 arranges the Equatorial cohort from east to west (fig. S1f). However, one has to be careful not to over interpret the patterns in higher PCs as these PCs explain very small fractions of variance (from 1.41% in PC 3 to 0.31% in PC 10).

##### Fixation indices

On chromosome arm 2L, pairwise F_ST_ values between geographic cohorts reveal a structure compatible with that observed in PCA (fig. S2e). A similar pattern among geographic cohorts is observed on the X chromosome, which also does not contain any common inversions (*19*). Interestingly, the X chromosome displays lower F_ST_ values among Equatorial cohorts than chromosome 2L, while the reverse is true among Southeastern, Central and South Mozambique cohorts. This is probably related to the relative difference in genetic diversity on the sex chromosomes and autosomes for the different cohorts (discussed below).

##### Doubletons

Doubletons are alleles that occur exactly twice in a dataset. They tend to represent rare variants in the population and are more informative for recent demographic events than higher frequency variants, but one major caveat is that not all doubletons represent true single origin variants, with PCR errors, sequencing errors, and convergent mutation all contributing to non-single origin doubletons. Within subset_3 we identified 9,811,404 doubletons and recorded in which cohort(s) they occurred (fig. S2f). 11.4% of all accessible sites have more than one alternative allele (i.e. triallelic or quadallelic), indicating that the infinite sites model does not hold true and we expect a considerable proportion of our observed doubletons do not have a single origin. However, single origin doubletons should be distributed in accordance with recent population diversity and connectivity, and doubletons caused by recurrent mutations and sequencing errors should be distributed evenly across pairs of cohorts, thus adding uniform noise to the doubleton sharing counts. When a cohort is representative of a spatially structured and localled panmictic population, we expect it shares more doubletons within itself than with other cohorts and this is indeed what we see for all cohorts. The individuals in Equatorial cohorts carry more doubletons than other cohorts, as is expected from the lower nucleotide diversity in the latter cohorts. The population structure revealed by doubleton sharing is similar to that observed in PCA and pairwise F_ST_. A subtle difference seems to be that doubleton sharing is stronger between the Equatorial than the Central cohorts, suggesting that the populations in the Equatorial region are more connected and have been mixing in the recent past compared to those in the Central region.

#### Genetic diversity

The majority of genetic diversity statistics presented here are calculated on subset_2 using the sc site filter (Methods). We explicitly state if and why a different sample subset, cohort definition or site filter was used.

##### Number of heterozygous sites

Counting the number of heterozygous sites per individual, we found that cohorts displayed a tight range of values, with some outliers with a lower or higher number of heterozygous sites than expected (fig. S2a). The high outliers were mostly individuals with median coverage below 20x, and we suspect that due to lower coverage, PCR or sequencing errors will occasionally be called as variants in these individuals (Methods). Restricting the plot to subset_2 individuals removes most of the high outliers (Fig. 1b). The lower outliers typically displayed elevated ROH counts and fractions, hinting at some recent inbreeding events, possibly due to bottlenecks (figs. S2b, S3d). The KE-N and MZ-M cohorts contained two and 19 males respectively, and since the average number of heterozygous sites in males was markedly lower than in females from the same cohort (2.532 M in males versus 2.791 M in females for KE-N, 2.290 M in males versus 2.345 M in females for MZ-M), most likely driven by the haploid X chromosome in males, the males were removed from the figure. Segregating inversions also contributed to the variation in the number of heterozygous sites per individual within the same cohort. When we excluded the genomic regions overlapping segregating inversions, that is sites which fall within the inversion boundaries (table S2, Tab “Genomic Coordinates”), the relative differences between individuals from the same cohort shrunk (data not shown).

##### Nucleotide diversity

The Equatorial cohort had higher nucleotide diversity than the Southeastern cohort, and this observation holds true for every chromosomal arm separately. However, there are differences between chromosomal arms, most notably between the X chromosome and autosomes. The neutral expectation is that nucleotide diversity on the X is 0.75 of the nucleotide diversity on autosomes, but this ratio can be affected by genetic drift, population bottlenecks, selection, mutation rate variability and recombination rate variability (*100*). We computed the nucleotide diversity ratio between the X chromosome and autosomes by computing the mean nucleotide diversity using 20 kb non-overlapping windows on geographic cohorts including individuals from subset_2 (table S2, fig. S2c). Interestingly, this ratio equals the neutral expectation for the Central cohorts, while for the Southeastern and South Mozambique cohorts it is lower, and for the Equatorial and North Ghana and South Benin cohorts it is higher. This suggests that different demographic or environmental processes are affecting the X to autosome diversity ratio in the Equatorial and Southeastern cohorts.

On each autosomal chromosome arm, the South Mozambique and the Central and Southeastern cohorts have Tajima’s D values close to zero while the Equatorial cohorts have values between -1 and -2, with South Benin and North Ghana interpolating in between. On the X chromosome, the Central and Southeastern cohorts have slightly positive values. Combined with the observation that for these cohorts nucleotide diversity is reduced on the X chromosome compared to autosomes, this implies that Watterson’s estimator (θ), which estimates mutation rates in populations, is even more strongly reduced on the X chromosome compared to the autosomes, which indicates a lack of rare alleles on the X and can be due to e.g. a recent selective sweep, balancing selection specifically affecting the X chromosome, or a sex-biased demographic event.

#### Differentiated populations

##### The South Benin cohort may be a new ecotype

Folonzo and Kiribina are two distinct chromosomal forms of *An. funestus* s.s. occurring sympatrically in Burkina Faso (*24*). The Kiribina form is characterised by its fixed homozygous standard karyotype for the 3Ra, 3Rb and 2Ra inversions. Because the Benin_Atlantique-Dept cohort (BJ) appears to be an outlier compared to its neighbouring populations (Figs. 1c, 2e, fig. S4e) and displays a similar karyotype to Kiribina (Fig. 2e, fig. S4e), we wanted to compare BJ to these known chromosomal forms from Burkina Faso. To this end, we aligned publicly available sequence data of 68 Kiribina and 86 Folonzo individuals (*23*) to the AFunGA1 reference genome, and followed the same procedure for variant calling and sample QC as already described. We performed PCAs on a combined dataset of Kiribina and Folonzo from Burkina Faso, as well as our Equatorial, South Benin and North Ghana cohorts (fig. S3b). The BJ samples diverge from all other *An. funestus* s.s. from the same geographical region to an even greater extent than Kiribina diverges from Folonzo. Based on existing literature we believe that Kiribina is an ecotype specific to a small region of Burkina Faso (*24*, *101–103*), therefore it is possible that the BJ samples are also representative of a new *An. funestus* s.s. ecotype.

Besides unexpected fixed inversion karyotypes based on neighbouring population karyotypes, Benin samples also show two regions of strong differentiation compared to all other cohorts, on chromosome arm 3R:36.5-37.1 Mb and on chromosome arm 2R:46.9-47.4 Mb (Fig. 1d, fig. S3c). We looked for non-synonymous amino acid changes in the genes within these regions, but did not find anything that was at high frequency in BJ and low frequency in other cohorts or vice versa (table S3). We did however find several synonymous SNPs with considerable frequency differences between BJ and other cohorts. Additionally, we saw many fixed variants in the intergenic region between two protein-coding genes annotated as ‘semaphorin-2A-like’, the region on chromosome arm 3R where the differentiation between BJ and other cohorts is strongest (fig. S10). Semaphorin-2A is a series of proteins that in *Drosophila melanogaster* are known to regulate transmembrane receptors, adult behaviour, motor neuron survival, and salivary gland positioning (*104*). We suspect that the peak of divergence is not driven by selection on non-synonymous SNPs within coding sequences, but rather on upstream or downstream regulatory regions or that there may be sequence in the genomes of BJ individuals that is not represented by the AfunGA reference genome, for example transposable elements.

*An. funestus* in Benin have been reported to be exceptionally resistant to DDT, pyrethroids and bendiocarb (*43*). Additionally, they have been found in abundance during the dry season and display an exceptional adaptive potential (*105*). We don’t know whether the regions of high divergence we report here are related to these notable attributes of the *An. funestus* populations in Benin, but it would be interesting to further investigate this. Given the potential role of transposable element insertions and structural variation in adaptation, future research would ideally use long-read sequencing to explore these populations.

##### The North Ghana population is bottlenecked and may also be an ecotype

The North Ghana (GH-N) cohort displays low genetic diversity in comparison to its geographically proximal neighbours (Fig. 1b, fig. S2a,c) and PC2 separates GH-N from all other cohorts (Fig. 1c). GH-N is also characterised by long runs of homozygosity (figs. S2b, S3d). All these signals are consistent with GH-N going through a recent population bottleneck.

Apart from reduced genetic diversity, the GH-N also displays increased divergence along part of the genome. Compared to its geographically closest neighbour, Ghana Ashanti-Region, as a representative of the Equatorial cohort, GH-N shows strong differentiation in the region of overlapping inversions on chromosome arm 2R and on the entire chromosome arm 3L (fig. S3e). The two cohorts have very different karyotype frequencies for the 2Ra and 2Rh inversions (fig. S4e), so the differentiation in this particular region is expected. Less expected is the elevated F_ST_ on the entirety of the 3L chromosome arm; in fact this differentiation is so strong, that the North Ghana cohort could not be confidently karyotyped for the 3La inversion (Fig. 2e, fig. S3f). Potentially the 3L arm experienced introgression from a closely related species before or shortly after the bottleneck. Beyond this, as mentioned in the main text, one individual collected in the GH-N region was genetically assigned to the Ghana Ashanti-Region cohort, suggesting that there may be sympatric diverged ecotypes in North Ghana. Together, these results indicate that North Ghana may harbour ecotypic variation in *An. funestus*.

#### Inversions

Large polymorphic inversions are common in the Gambiae Complex and the Funestus Subgroup (*106*, *107*). Because recombination between the two inversion orientations is suppressed in heterozygous individuals, inversions can link beneficial adaptations in several genes, effectively acting as ‘supergenes’ (*108*). The lengths of the inversions discussed here range from 8.4 to 19.7 Mb, in total accounting for more than 30% of the nuclear genome. Polymorphic inversions in *An. gambiae* and *An. funestus* have been linked to ecological and behavioural adaptation, in particular aridity tolerance (*31*, *109–115*).

##### Inversion breakpoint identification

We analysed and compared the two publicly available chromosome-level genome assemblies for *An. funestus*: AfunGA1 (GCA_943734845.1) generated from a single individual representing a wild population in Gabon (*18*) and AfunF3 (GCA_003951495.1), which used pooled individuals from the FUMOZ colony originating from samples collected in Mozambique (*116*).

Centromeres of AfunGA1 were preliminarily defined as >100 kbp regions of highly homogenous tandem repeats surrounded by >1 Mbp repeat rich regions of pericentric heterochromatin (table S2, Tab “Genomic Coordinates”). Base-pair resolution coordinates were identified from a combination of AfunGA1 genome self homology inferred in StainedGlass (*117*) v0.5 and visualised in HiGlass (*117*) v1.11.6 for preliminary region detection coupled with ULTRA (*118*) v0.99.17 for tandem repeat units identification and precise repeat regions annotation. For predicting heterochromatin regions we combined StainedGlass results with transposable elements annotations from Earl Grey (*99*) v2.1. Heterochromatin boundaries were arbitrarily identified at 10 kbp resolution based on elevation in transposable element density compared to neighbouring euchromatic regions.

Inversion breakpoints for 2Rh, 3Ra, 3Rb and 3La were estimated using SyRI (*119*) v1.6.3 based on minimap2 (*120*) v2.24 whole-genome alignments between AfunGA1 and AfunF3 assemblies. Breakpoint coordinates are given in base-pair resolution as the ends of aligned inverted segments (table S2, Tab “Genomic Coordinates”). D-GENIES (*121*) v1.5.0 genome alignment dot plots based on the minimap2 alignments were used to cross-check the SyRI result and investigate inversion breakpoint regions in more detail. In particular, we found that the right breakpoint of 2Rh as well as the left breakpoint of 3Ra contained small nested inversions. The right breakpoint of 3La fell within a potential mis-assembly in AfunF3 resulting in a translocation combined with a duplication. As a result, there are several candidate locations of the breakpoint based on the alignment and they span a range of 250kb.

The breakpoints of inversion 2Ra and 2Rt could not be estimated with SyRI because AfunGA1 and AfunF3 carry the same orientation for these inversions. Instead, we tested an F_ST_-based method to estimate breakpoint coordinates for segregating inversions within our dataset. We computed F_ST_ values of single variants by comparing homozygous standard (2R+^a^/2R+^a^) individuals to homozygous inverted individuals (2Ra/2Ra). We noticed that near the inversion breakpoints there is a very sudden increase in the number of variants with high F_ST_. We also computed the F_ST_ values in sliding windows of 6,000 bp, with a step-size of 1,000 bp, and report the inversion breakpoint to be the window-centre of the local maximum in the sliding-window mean near the sudden increase of F_ST_ values (table S2, Tab “Genomic Coordinates”, fig. S1c). For the four inversions that had breakpoints determined by SyRI, the breakpoints reported by the described F_ST_ method were on average 1.8 kb and at most 4.2 kb off. We followed the same procedure to estimate the breakpoints of other uncertain inversion breakpoints, such as the 2Ra inversion, which is split into two parts when aligned to AfunGA1, because it overlaps with the 2Rh inversion, the 2Rt inversion, and the right breakpoint of 3La, because its inversion breakpoint could not be unambiguously determined by the comparing reference genomes due to discontinuities in the reference genome alignment. The coordinates we report and use are those inferred from comparing reference genomes (2Rh, 3Ra, 3Rb, left breakpoint of 3La) and those inferred by the F_ST_ method (2Ra, 2Rt, right breakpoint of 3La) (table S2, Tab “Genomic Coordinates”).

##### Identifying inversions by literature comparison

A total of 17 segregating inversions in *An. funestus* have been previously described (*19*). Because all six inversions discussed in this study occur in multiple cohorts, meaning they are reasonably common, they can likely be linked to these known inversions. For inversions 3Ra and 3Rb, the genomic coordinates of breakpoints are known in AfunF3 (*122*), and are very close to the inversion breakpoints inferred by comparing AfunF3 and AfunGA1. Similarly, the genomic coordinates for 2Ra (*122*), when lifted over from AfunF3 to AfunGA1, are close to those inferred by the F_ST_ method. We used the microsatellite photomap (*19*) to identify candidate names for the remaining inversions based on approximate size and position. There are three known segregating inversions on chromosome arm 3L, and only 3La occurs throughout sub-Saharan Africa (*19*, *115*, *123*). Moreover, we mapped microsatellite FUN K (*124*) to the AfunGA1 reference genome, and it falls inside the observed inversion region, which is consistent with 3La and inconsistent with any other described inversions on this arm. On 2R we observe the 2Ra inversion and two other inversions, one overlapping 2Ra from the left and one overlapping 2Ra from the right. In either case, 2Ra does not contain all of the other inversion, nor does the other inversion contain all of 2Ra. The microsatellite map (*19*) shows that only 2Rt and 2Rh meet these criteria for the left and right respectively. Moreover the primers for microsatellite AFND32 (*125*) map to the region inside the 2Rh inversion and outside the 2Ra inversion, further confirming it is the 2Rh inversion overlapping 2Ra from the right.

##### In silico karyotyping

Our sliding window PCA showed four regions where the most prominent structure groups the samples by their inversion karyotype (one on 2R, two on 3R and one on 3L, Fig. 1d). We performed *in silico* inversion karyotyping by defining two threshold PC1 values that separate the three karyotypic states (homozygous inverted, heterozygous, homozygous standard) and assigning karyotypes accordingly. Although the inversion karyotype is the most prominent structure in the inversion regions, existing population structure still affects the position of samples on PC1 and hence we set separate thresholds for different PCA cohorts (Fig. 2a, fig. S5d). For the 3La inversion we noticed a decay in karyotype separation towards the middle of the inversion, so we performed *in silico* karyotyping separately for the inversion regions near the left and the right inversion breakpoint, and found that all samples had concordant karyotype results for both regions (Fig. 2c).

On arm 2R we noticed a complex inversion region, corresponding to the known overlapping inversions 2Ra, 2Rh and 2Rt. The overlap of 2Ra and 2Rh, which are more wide-spread than 2Rt in our dataset, results in three distinct inversion-driven patterns: only 2Ra is segregating, only 2Rh is segregating, 2Ra and 2Rh overlap (fig. S4, section overlapping 2R inversions below). We performed *in silico* karyotyping for these three regions separately: for the region where only 2Ra is segregating we allowed for three karyotypes (2R+/+, 2Ra/+, 2Ra/a), for the region where only 2Rh is segregating we initially allowed for three karyotypes (2Ra/a, 2Ra/+, 2R+/+), and for the region where 2Rh and 2Ra overlap we allowed for six karyotypes (2R+/+, 2Ra/+, 2Ra/a, 2R+/h, 2Ra/h, 2Rh/h) (fig. S4d). We karotyped the samples by setting one set of two PC1 threshold values in the region where only 2Ra segregates and a different set of two PC1 threshold values in the region where only 2Rh segregates, again allowing for different thresholds for the different PCA cohorts, due to the compound signal of inversions and geographic population structure. We then checked whether the combined 2Ra and 2Rh karyotypes were consistent with the sample trajectories in the region where these inversions overlap.

We discovered two additional ‘intermediate’ karyotypes in the region where 2Ra, but not 2Rh, is segregating, occuring only in DRC_Haut-Uele and Cameroon. These two intermediate states turned out to be driven by heterozygous individuals for the relatively rare 2Rt inversion (*19*) (fig. S4cde).

The three different trajectories in the sliding window PCA, or the three different clusters in a two-dimensional PCA, correspond to the three different inversion karyotypes. To find out to which homozygous inversion orientation the top and bottom trajectory corresponds, we incorporated the reference genome AfunGA1 into our sliding window PCA as an individual with homozygous reference calls on the entire genome. With the assumption that the karyotype of AfunF3 is 2R+^a^+^h^ 3R+^a^+^b^ 3L+^a^ (Igor Sharakhov, pers.comm.) and the karyotype of AfunGA1 is 2R+^a^h 3Rab 3La (*18*), this enabled us to assign standard and inverted orientations to the homozygous bands across our samples.

As a confirmation, and to karyotype the samples from North Ghana that were not incorporated in the sliding window PCA, we performed two-dimensional PCAs using variants from the entire inversion region. Samples indeed clustered according to the karyotype assigned from the sliding window PCA. The North Ghana samples fell within a single karyotype cluster for the 2Rt, 2Ra, 2Rh and 3Rb inversions. The 3Ra inversion is segregating in the North Ghana cohort, and we found that some samples fell in the 3Ra/a cluster and the others in the 3R+/a cluster. The North Ghana cohort is strongly diverged on the 3L chromosome arm (see section North Ghana bottleneck above), including on the 3La inversion region, and could therefore not be confidently karyotyped for this inversion.

##### Hardy-Weinberg Equilibrium

An observed violation of Hardy-Weinberg equilibrium (HWE) due to a lack of heterokaryotypes in several villages in Burkina Faso in the late 1990s led to the proposal of two sympatric *An. funestus* populations that were reproductively isolated (*24*). These two populations, named Folonzo and Kiribina, were characterised by a deterministic algorithm taking into account the 2Rs, 2Ra, 3Ra and 3Rb karyotypes (*24*). The 2Rs inversion was assumed to be only segregating in Kiribina, while the other three inversions were assumed to be segregating in Folonzo and found only at very low frequencies in Kiribina. The algorithm was designed to ensure both populations satisfied HWE.

For our sample set, we assessed HWE for each inversion karyotype within every geographic cohort using Pearson’s χ^2^ statistic with one degree of freedom. Here we report the cohorts and inversions violating HWE at a significance level of 0.05, i.e. with χ^2^ ≥ 3.84. Violation of HWE is a sign that the cohort does not represent a single panmictic population and might warrant further investigation to understand the local population structure and the impact that might have on the success of vector control measures.

The Kenya Nyanza Province geographical cohort is out of HWE for 3Ra (χ^2^ = 33.42), 3Rb (χ^2^ = 6.55) and 3La (χ^2^ = 6.75). This cohort contains samples from five different collection locations (table S1). Ahero, the northernmost location, has very dissimilar inversion frequencies from the other locations. However, we suspect that this is driven by seasonality or a temporal population shift rather than by geography, as samples from Ahero were collected in June 2014 compared to samples from the other four locations which were collected in October 2016. The Nyanza Province has two wet seasons, one from March to June and a second from September to December (*126*). When we partition the samples from Kenya Nyanza Province by month of collection, HWE is satisfied within both groups for all inversions.

For the rare 2Rt inversion, which we only detected in cohorts Cameroon_Adamawa and DRC_Haut-Uele, all observed individuals carrying this inversion are heterozygous (2Rt/+), in which case HWE is satisfied in both cohorts.

##### Associations between inversions

We tested for associations between inversions within PCA cohorts using Huff and Rogers *r* statistic to detect linkage disequilibrium (*127*) (table S2). Note that North Ghana and South Benin are omitted from the table, because they each have only one segregating inversion. We observe a negative correlation between 2Ra and 2Rh, which is not surprising because they overlap and hence cannot occur on the same chromosome. 2Ra and 2Rt also overlap, but because we have very few samples with the 2Rt inversion, we do not have the power to detect a strong negative correlation. We find a positive association between all pairs of the three inversions on the 3RL chromosome, suggesting that perhaps there are interactions between these inversions resulting in an additional advantage when carrying multiple of these inversions.

##### Double recombinants

While we typically observe three trajectories within inversion regions (corresponding to the three inversion karyotypes) in the sliding window PCA, in some places we see that a single sample line joins a different trajectory for parts of the inversion region (Fig. 2b), suggesting that alleles carried by the individuals corresponding to these lines are the product of double recombination.

We found double recombinant individuals for the 3Rb and 3La inversions, the two longest segregating inversions in the dataset. For each putative double recombinant sample, we performed a 2D PCA on biallelic variants with minor allele frequency ≥0.02, using the sc filter, from a 1 Mbp region centred on the trajectory change, to confirm that the individual indeed carries variants consistent with a locally different karyotype (fig. S5ef). Not all putative double recombinant individuals cleanly clustered with a different karyotype in the 2D PCA performed on this inner-inversion breakpoint; it is possible that the trajectory change in these individuals was caused by noise rather than recombination, or that the double crossover breakpoints occur so close together that a 1Mb window does not provide sufficient resolution.

Possibly, because of its size, age, or historic frequencies, the 3La inversion has experienced increased genetic exchange between the inversion orientations. This could be the cause of the pattern we observe for 3La in the sliding window PCA (Fig. 2c, fig. S5b), where the three karyotypes are clearly separated near the breakpoints, but the separation decays towards the centre of the inversion.

At 30.5 Mb on chromosome arm 3R, we observe a double recombinant from MZ-M, as a line from the top trajectory (3R+/+) briefly joining the middle trajectory (3Rb/+), meaning that this homozygous standard individual locally exhibits a heterozygous karyotype (Fig. 2b, fig. S5e). At the same place in the genome, we observe that the single homozygous standard individual from GA, as well as all heterozygous individuals from GA, also show a dip in the sliding window PCA toward the lower (3Rb/b) trajectory (Fig. 2b). We cannot pin the signal down to a particular gene, but of potential interest are four genes characterised as ‘troponin C’ (AFUN2_001408, AFUN2_008892, AFUN2_009192, and AFUN2_012458). As part of the troponin complex, troponin C is crucial for muscle contraction (*128*). In the moths *Mythimna separata*, it has been shown that the expression levels Troponin C were upregulated in response to the botanical insecticide wilforine (*129*). We did not find any non-synonymous or synonymous SNPs in coding regions that were at different frequency in heterozygotes in Gabon compared to heterozygotes from other cohorts. However, because the response to wilforine was upregulation of Troponin C, we expect that any adaptation is likely driven by regulatory elements rather than SNPs within coding regions. We have not found reliable records of wilforine use in the region where these samples were collected in Gabon, but the genomic signal is consistent with a local selective pressure favouring the alleles found on the 3Rb orientation.

It is widely accepted that polymorphic inversions result in suppressed recombination rate in heterozygous individuals (*130*). Some efforts have been made to quantify the extent of recombination suppression, showing in *Drosophila* flies that double recombinants do occur, but are relatively rare (*130*). The sliding window PCA allows for easy identification of candidate double recombinants and the signal we observe in the GA cohort suggests that double recombinant alleles might be positively selected if they contain adaptive variation.

##### Overlapping 2R inversions

The 2R chromosome arm contains several overlapping inversions (*19*), three of which (2Ra, 2Rh, 2Rt) we found segregating in our dataset. We first focused on 2Ra and 2Rh, since these are segregating in the majority of our cohorts. For simplicity, throughout this section we refer to the telomeric end of the 2R chromosomal arm as the “left side” and the centromeric end as the “right side”.

The 2Ra and 2Rh inversion regions partially overlap, with sub-regions from left to right where only 2Ra segregates, both 2Ra and 2Rh segregate, and only 2Rh segregates. The AFunGA1 reference carries the 2Rh inversion, and alignments to this reference genome rearranges the order of the three regions: the 2Ra and 2Rh overlap is now on the right and the 2Ra region is split into two unconnected parts (fig. S4d).

Because 2Ra and 2Rh overlap, a chromosome cannot be inverted for both at the same time, so we have three possible alleles: the standard orientation (2R+), the 2Ra inversion (2Ra) and the 2Rh inversion (2Rh). This results in six possible combined karyotypes for the two inversions: 2R+/+, 2Ra/+, 2Ra/a, 2R+/h, 2Ra/h and 2Rh/h. In the sliding window PCA (fig. S4b,d), there are three states in the left region (where only 2Ra is segregating), corresponding from top to bottom to homozygous standard, heterozygous and homozygous inverted for 2Ra. In the middle region (where only 2Rh is segregating) there are also three states, corresponding from top to bottom to homozygous standard, heterozygous and homozygous inverted for 2Rh. In the right region (where 2Ra and 2Rh overlap), there are six states, corresponding to the six combined karyotypes in the order listed above.

In the region where 2Ra is segregating and 2Rh is not segregating, the sliding window PCA exhibits two additional states in between the three states corresponding to 2Ra karyotypes (fig. S4c,d). These states contain two individuals from Cameroon_Adamawa and eleven individuals from DRC_Haut-Uele, and are driven by the 2Rt inversion. However, because this inversion is only carried by a few individuals and is most likely only present in a heterozygous state (fig. S4a), its signal is not strong enough to drive the windowed PCA on the entire 2Rt region when all samples are included. Nevertheless, in regions where the PCA is already driven by the 2Ra inversion, i.e. the inversion signal is stronger than the geographic signal, we can also observe the additional structure caused by the 2Rt inversion.

Because 2Rt and 2Ra overlap, the three possible alleles with respect to these inversions are 2R+, 2Rt and 2Ra. This results in six possible combined karyotypes: 2R+/+, 2R+/t, 2Ra/+, 2Ra/t, 2Ra/a, and 2Rt/t. As mentioned previously, 2Rt is fairly rare (fig. S4e), and we do not observe any homozygous 2Rt/t samples in our current dataset. We do, however, see all five of the other expected states for the combined 2Rt and 2Ra karyotypes (fig. S4c,d). We observe a total of 11 combined karyotypes of the three inversions segregating on chromosome arm 2R, each with a distinct trajectory in the sliding window PCA plot. Due to limitations of our short Illumina generated reads, we cannot tell the difference between the 2R+/ht and 2Rh/t karyotype. However, considering we do observe individuals that are heterozygous for each of 2Rt, 2Ra and 2Rh, there must be an allele containing both the 2Rt and 2Rh inversion. Whether both regions are inverted separately or whether the combined region is inverted can only be resolved by looking at long read data.

##### Heterozygosity of inversion orientations

For all reported inversions we compared the mean heterozygosity within each inversion region for homokaryotypic individuals (because we only have heterozygous and homozygous standard karyotypes for 2Rt, this inversion was not considered in this analysis). The phased haplotypes contain many switch errors throughout the inversion regions, so we conducted this analysis on genotypes rather than haplotypes and disregarded individuals that are heterozygous for the inversion. To account for population structure we performed this analysis separately for each PCA cohort with at least ten individuals per homokaryotype. We computed mean heterozygosity in non-overlapping windows of 100 kbp accessible sites with the sc filter. All cohorts with sufficient homokaryotypic individuals showed a similar pattern, only the Equatorial cohort is shown in fig. S5c. For 2Rh, the homozygous inverted karyotype has lower mean heterozygosity than the homozygous standard karyotype. Such a difference was not observed for the other inversions. While we would expect the derived orientation to have lower heterozygosity than the ancestral orientation, there are many factors to take into consideration that could explain the contrasting patterns between these inversions, such as the demographic history of the population where the inversions are segregating, the historic allele frequency trajectory, natural selection, the strength of recombination suppression and the age of the inversion (*131*).

#### Recent selection and insecticide resistance

##### Detecting selective sweeps with H12

In order to detect potential hard and soft selective sweeps, we ran the H12 statistic (*35*) on phased haplotype data from all geographic cohorts. The window sizes used in the H12 analyses have to be calibrated separately for each cohort, because different demographic histories and different levels of genetic variation affect the expected values of the H12 statistic (*35*). We tested window sizes ranging from 100 to 4,000 SNPs per window and selected for each cohort and each chromosome the smallest window where 95% of windows have mean H12 value below 0.1 (table S2). We set the threshold to classify sites under selection as peaks with H12 values > 0.4. The high levels of homozygosity in the GH-N cohort resulted in overall high H12 values, where it was difficult to distinguish signals of relatedness from potential selective sweeps, so we excluded GH-N from the H12 analysis and haplotype tree generation.

##### Sites under selection

From the H12 scan, we identified four loci (*Gste2*, *Gaba, rp1*, *Cyp9k1*) that are under selection in multiple cohorts and two additional loci under selection in a single cohort (*Vgsc*, X: 13.7Mbp). We assessed SNP frequencies in all genes within a 0.2 Mbp region centred on the height of the H12 peak to determine which genes were likely under selection; in most cases we found genes that were previously described as (candidate) selection sites. We constructed haplotype trees for the genes putatively under selection. In most cases, the structure of the haplotype trees was stable over different genomic regions in the vicinity of the putative genes under selection. However, for very small genomic regions containing few variants, the trees might differ considerably from region to region (e.g. for single *Gste* genes). Also for the peak at 13.7 Mbp on the X chromosome, the haplotype trees differ depending on the genomic region used, probably because there is very low accessibility in this region in general. Here we discuss the candidate variants under selection, their geographic spread and the haplotypic backgrounds on which they occur.

##### Gste2

The H12 peak falls within a region containing seven *Gste* genes. Some individuals in our dataset bear the known L119F variant in the *Gste2* gene (AFUN2_012779). This variant is only observed in the non-canonical transcript 2 of the gene. Moreover, this gene is annotated as “glutathione S-transferase 1-like” in AFunGA1, rather than as *Gste2* or *Gste2-*like as expected. These are probably annotation issues with the relatively new reference assembly. On top of the known variant, we also observe another variant in the same amino acid, L119V, found at low frequencies in east Africa. But as it is found scattered throughout the haplotype tree, it does not show any signal consistent with a recent selective sweep.

Within the other six *Gste* genes we have noticed the presence of other non-synonymous SNPs present at similar frequencies to L119F in the same populations (G26D in AFUN2_000742, A315T in AFUN2_012863). More research is needed to check whether these are merely neutral on the same haplotypic background as L119F, or whether these additional amino acid changes across multiple *Gste* genes convey a selective advantage to carriers of the L119F mutation.

##### Gaba (rdl)

It has been shown in *An. gambiae* that the A296S and A296G mutations in the *Gaba* gene changes the shape of the dieldrin binding site, thereby conferring resistance to this insecticide (*34*). In the Gambiae Complex, the rdl locus lies within the 2La inversion region and the occurrence of either resistance mutation is strongly associated with the 2La karyotype (*34*). Although in *An. funestus* the *Gaba* gene falls outside the 3Ra inversion region, the A296S variant is significantly correlated with the 3Ra inversion karyotype in geographic cohorts where both are variable and A296S is at frequency ≥0.02 (GH-N, GH-A, NG, CM, MW; χ^2^-test with two degrees of freedom, values 10.922 - 30.167 [p-values 0.0042 - 3×10^-7^]), except for CD-K (χ^2^ = 5.198 [p = 0.074]). Given these patterns, it may be that the rdl A296S mutation confers increased resistance when linked to variants in the associated inversions.

In the Gambiae Complex, another non-synonymous variant in the *Gaba* gene, T345S, was reported to be in strong linkage disequilibrium with the A296S variant (*34*). The authors hypothesise that the codon 345 mutations could compensate for the fitness costs induced by the A296S mutation in absence of dieldrin, and thereby help explain why the A296S variant is still found decades after dieldrin is thought to have ceased being used. In *An. funestus* we also observe a correlation between the A296S and T345S variants (linkage disequilibrium [LD] Rogers and Huff *r* (*127*) in the six cohorts where both variants are observed: 0.494 (GH-N), 0.641 (GA), 0.716 (BJ), 0.733 (UG), 0.930 (MZ-M), 0.930 (TZ)), which would be consistent with the compensatory fitness effect of the latter. However, T345S is at lower frequency than A296S in all cohorts and there are six cohorts where A296S is found but not T345S, so the correlation is only observed on regional scales.

The non-synonymous V327I mutation has been reported to co-occur with the A296S mutation in *An. funestus* individuals from Burkina Faso and Cameroon (*38*). Here, we find only two individuals carrying the V327I mutation, both in a heterozygous state. These samples, VBS24121 and VBS24128, from South Mozambique, are also heterozygous for the A296S mutation. When this variant was first described, it was found at intermediate frequencies in central Africa and absent in cohorts from eastern and southern Africa (*38*), while now we find it at very low frequency and only in southeast Africa.

##### rp1

A strong H12 peak on chromosome 2R is present across all geographic cohorts, centred on a well-studied cluster of Cytochrome 6 P450 (*Cyp6p*) metabolic genes known as the “resistance to pyrethroid” (*rp1*) locus (*132*) (Fig. 3a, table S3). Haplotype clustering on the whole locus results in a pattern where the haplotypes of each individual are split between two clusters possibly indicative of copy number variants (CNVs). CNVs play an important role in conferring metabolic insecticide resistance at the *rp1* locus in *An. gambiae* (*133*) therefore we examined this locus for CNVs. Exploration of coverage and discordant read mapping in the region (see Methods) identified nine independent CNVs, eight of which overlapped with Cyp450 genes (fig. S7a). Haplotype clustering outside of the region containing the detected CNVs, results in haplotype clusters tightly correlated with geography (fig. S7a,b). In West Africa, most clusters were associated with the presence of a single CNV. We could not detect any CNVs explaining sweeps in East African populations, but we did find the previously described 6.5 kb insertion (*134*) associated with the Southeastern and South Mozambique cohorts.

*rp1* is a complicated locus, where coding and regulatory SNPs, CNVs, and structural variants might all be undergoing selection(*15*, *134–137*). Regions like this motivate the generation of long read data from individuals and populations across the continent, in order to comprehensively study evolution and selection.

##### Cyp9k1

The H12 signal on the X chromosome present in six cohorts is centred on a single Cytochrome P450 called *Cyp9k1*, shown to contribute to metabolising both type I and II pyrethroids in *An. funestus* (*138, 139*) (Fig. 3a, table S3). Individuals from CF, CD-H, UG, KE-N, and KE-W share a haplotype carrying the G454A mutation that increases the catalytic efficiency of metabolising pyrethroids (*139*) (fig. S7c). Another swept haplotype, found in NG and GH-A, bears the mutation V327I, which has previously been reported at high frequency in Ghana, but its phenotypic effects have not yet been assessed (*139*).

##### Vgsc

The voltage gated sodium channel (*vgsc)* is the target of pyrethroid-based insecticides and DDT. Mutations at key sites in the *vgsc* gene cause target site resistance, referred to as *knock-down resistance (kdr)*, notably the L995F mutation in *An. gambiae* (*56, 140, 141*). While it was previously believed that *An. funestus* does not carry the *kdr* mutation, it has recently been shown that the L976F mutation (orthologous to the L995F mutation in *An. gambiae*) is nearly fixed in a Tanzanian population (*39*). Mosquitoes carrying this mutation were phenotyped and found to be resistant to DDT, which has been banned for use in this area since 2008. The high frequency of this mutation conferring DDT resistance in one region of Tanzania has likely occurred due a long term leaking DDT stockpile (*39*). The samples from our TZ geographic cohort show a sweep centred on the *Vgsc* gene in Tanzania containing the L976F *kdr* mutation.

##### X: 13.7Mbp

While the CM cohort has H12 > 0.4, several other cohorts have elevated H12 values at this locus that don’t pass the 0.4 threshold. Accessibility in this region is poor. The region contains one gene previously implicated in insecticide resistance in *An. gambiae* (AFUN2_000796; eye-specific diacylglycerol kinase) (*142*), however the H12 peak falls in a region between this gene and some smaller genes with no known insecticide resistance in other species. We do not see any relevant SNPs associated with CM in all genes spanning this 0.2 Mbp region (table S3), so possibly selection is acting on CNVs or regulatory elements.

##### Signals of selection in historic specimens

We explored whether any known insecticide resistance conferring mutations were present in historic specimens. We observe the G454A mutation in *Cyp9k1* in 13 out of 18 historic samples from 1936 Tanzania, with 12 samples showing a heterozygous state and one low coverage sample with a single sequence with the mutant SNP (table S4). This mutation is common across Africa but appears to only be associated with a swept haplotype in Eastern Africa (fig. S8a). The G454A sweeping haplotype identified here is likely to be the same haplotype that was recently found to be associated with pyrethroid resistance in the species in Eastern Africa and spreading west rapidly (*139*). The presence of the G454A mutation predating widespread insecticide usage suggests that the mutation likely has other important fitness benefits unrelated to resistance.

As discussed in the main text, the rdl A296S mutation in *Gaba* was detected in mosquitoes collected in west Africa in the 1960s. The first published records of insecticide resistance in *An funestus* that we were able to find date from 1957 for DDT (*44*) and 1964 for dieldrin (*143*). Considering this fact and the fact that DDT was in widespread usage from the late 1940s onwards (*40*, *144*), it was surprising to find no evidence of the L119F mutation in the *Gste2* gene or the *kdr* mutation in the *Vgsc* gene, which both confer DDT resistance (*25*, *140*, *141*)

#### Gene drive

##### Target identification

We took a whole genome approach at identifying potential gene drive targets in *An. funestus*. This approach comes with several caveats: it does not take into account gene function, it only looks for 20bp sequences that fall entirely within coding sequence, and it relies on the accuracy of gene annotation in the reference genome. However, in reality, gene drive target sites can range in length from 17 to 24 bp (*145*), they could be on an intron-exon boundary, instead of falling entirely within coding sequence, and potentially some exons are missing from the current AFunGA1 gene annotation (*74*) (version 65), as demonstrated by investigation of the doublesex target site (see below). Moreover, this approach does not take into account gene function, while this is important for suppression gene drives and to lesser extent modification gene drive (*17*).

We settled for identifying 20bp conserved sequences within coding sequence ending with the -GG protospacer adjacent motif (PAM) or, for the reverse complement, starting with CC-, where ‘-’ can be any nucleotide (and does not even need to be conserved within the population). These specifications are the same that were used to identify gene drive targets in Ag1000G phase 1 (*16*), so we can compare results between *An. funestus* and *An. gambiae* and *An. coluzzii*.

The choice of site filter has a considerable effect on the number of available target sites (table S2) and because this analysis is restricted to coding sequence, which should be reasonably conserved within the species, we conducted this analysis without a site filter. Previous studies showed that guide RNAs tolerate a certain level of mismatch (*146*), and we hope that by stringently selecting candidate target sites based on our dataset we minimise the risk of reduced detection efficacy due to unsampled variation that exists in natural populations.

We report the number of candidate target sites and their associated genes for *An. funestus* and *An. gambiae/An. coluzzii* with and without species-specific site filters (table S2, Tab gene drive targets). The ‘reference’ rows refer to sites identified when taking only the corresponding species reference into account (AfunGA1 and AgamP4). The ‘variation’ rows refer to targets where all samples in the datasets are homozygous for the reference genome at the required positions. For the rows titled ‘non overlapping’ we reduced the list of candidate targets to only non overlapping positions, such that they could in theory be simultaneously targeted.

##### Doublesex gene drive

A promising suppression drive in *An. gambiae* targets a highly conserved sequence located at the boundary of the female-specific exon in the *doublesex* (*dsx*) gene (*49*). The gene drive disrupts the formation of functional female splice transcripts, and results in sterility in females homozygous for the mutant allele, while homozygous males and heterozygous males and females have normal fertility levels. The *dsx* gene drive has successfully spread in both small laboratory cages (*49*) and large indoor cages incorporating more realistic vector ecology elements (*50*).

The 23 bp gRNA target site is conserved across at least five species in the Gambiae Complex (*49*) and there is only one nucleotide difference between *An. gambiae* and *An. funestus* (Fig. 4b). The high degree of conservation across species suggests that the target site does not allow for much variation. Natural variation in the population or newly arisen variation during repair of the double stranded break induced by the gene drive might lead to resistance to gene drive. We assessed the variation at the target sites in the wild populations represented in our dataset and in comparison to the available population genomic data for the Gambiae Complex.

In all *An. funestus* from our dataset (subset_1), we observe only one variant (Variant 1 in Fig. 4b), namely C>T at the fourth position, resulting in the same sequence as the AgamP4 reference genome. This variant is found at very low frequencies in two cohorts (CD-K: 1.5%, KE-N: 0.8% allele frequency). Among all *An. gambiae, An. coluzzii* and *An. arabiensis* mosquitoes from Ag1000G phase 3 of the MalariaGEN Vector Observatory (*72*) (3081 individuals total) we find two segregating variants in the target region. Variant 2 (Fig. 4b) at the nineteenth position is found in four cohorts and in both *An. gambiae* and *An. coluzzii* (AO-LUA_colu_2009: 25.9%, CD-NU_gamb_2015: 7.2%, GA-1_gamb_2000: 5.8%, CM-ES_gamb_2009: 1.5% allele frequency). Variant 3 (Fig. 4b) is found in heterozygous state in only one individual, and might indeed be a rare variant or a sequencing error (BF-09_gamb_2012: 0.5% allele frequency). The levels of natural variation at the gRNA target site of the *dsx* gene drive seem to be lower in *An. funestus* than in *An. gambiae* and *An. coluzzii*, so we are hopeful that the same approach can be translated to be used in *An. funestus*.

